# Mapping the immune landscape in small cell lung cancer unveils a distinct tumor-reactive CD8+ T cell molecular signature

**DOI:** 10.64898/2026.06.29.735200

**Authors:** Krutika Khinvasara, Elif Diken, Jennifer Gerbracht, Enes Huduti, Joshua D’Rozario, Tana Omokoko, Sebastian Newrzela, Özlem Akilli, Franziska Lang, Barbara Schrörs, Katja Höpker, Eliana Stanganello, Maik Schork, Alessandra Gargano, Ahmad Al Alwash, Jan-Phillip Weber, Julie George, Roman K Thomas, Ayline Kübler, Mustafa Diken, Uğur Şahin, Laura Kolb

## Abstract

Small cell lung cancer (SCLC) is a highly aggressive malignancy with limited therapeutic advances. Unlike many other cancers, its immune landscape, particularly immune competence and T cell recognition, remains poorly characterized. Here, we generate a single-cell transcriptome atlas of the SCLC immune microenvironment with paired T cell receptor (TCR) sequencing. By linking T cell states with clonality and a multilayered functional screening, we identify 6 tumor-reactive TCRs that recognize and eradicate autologous SCLC cell lines. We delineate a novel SCLC-reactive CD8+ T cell signature (SCLC_TR), enabling the identification of 47 further SCLC-reactive TCRs. The SCLC_TR signature performs extremely well in pancreatic ductal adenocarcinoma (PDAC), another immune-cold tumor indication, and, most strikingly, patients with elevated SCLC_TR signature scores exhibited significantly improved survival, underlining its prognostic potential. Comparative cell-cell interaction analyses implicate several immunosuppressive mechanisms, with myeloid cells and CD4⁺ regulatory T cells possibly acting as counterbalances to effector T cell activity in SCLC. In summary, our study challenges the prevailing notion of SCLC as an immune-cold tumor type by providing direct evidence of tumor-reactive T cell responses and introduces the SCLC_TR signature as a tool to identify tumor-specific T cells and their microenvironmental restraints and escape mechanisms, ultimately shaping next-generation immunotherapeutic strategies.

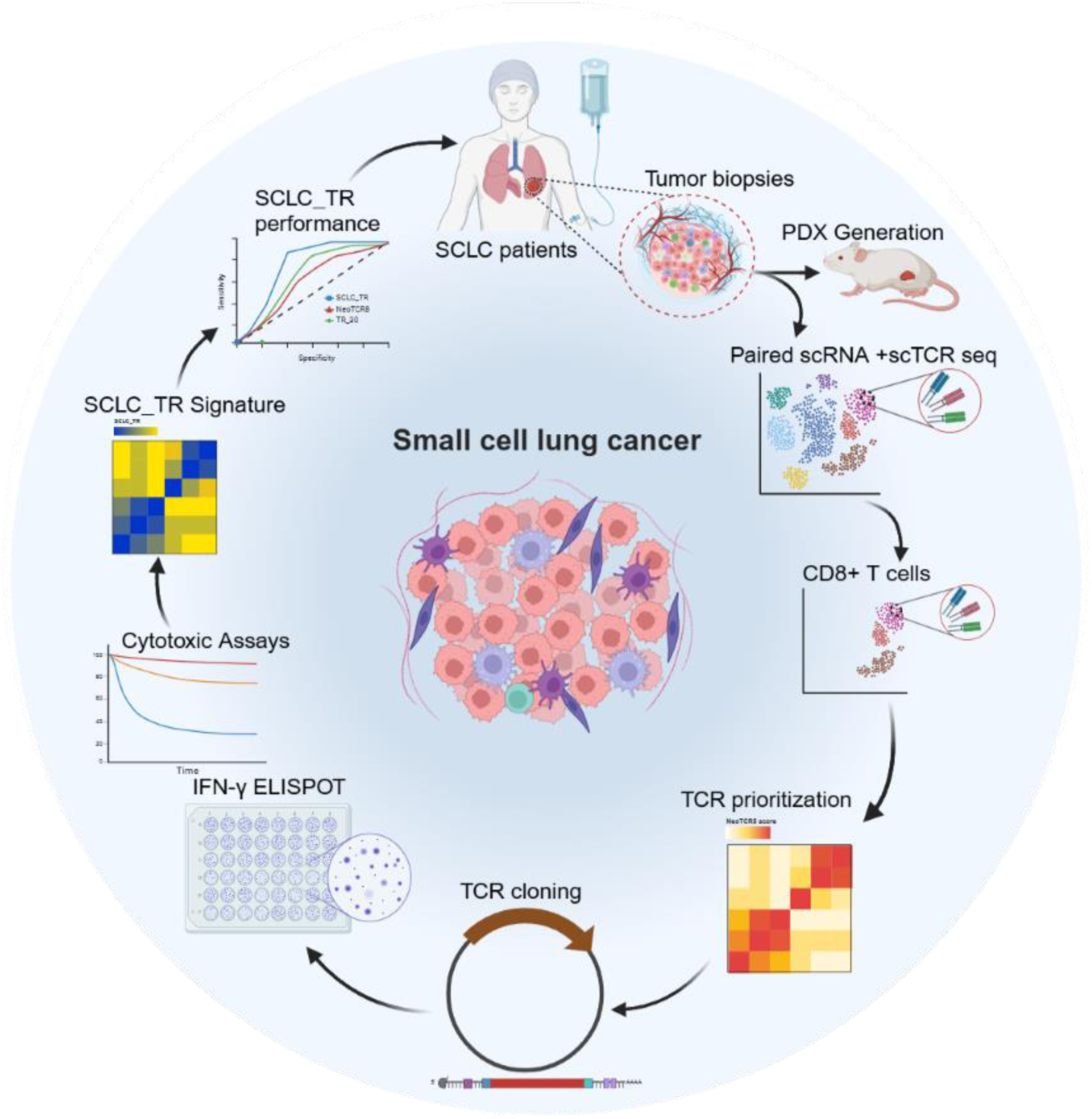

## Introduction

Lung cancer is the leading cause of cancer-related death worldwide, and small cell lung cancer (SCLC) represents its most aggressive subtype, accounting for approximately 15% of all lung cancer diagnoses.^1^ Characterized by rapid growth, early metastasis, and poor prognosis, SCLC has a 5-year overall survival rate of approximately 7%, with markedly lower survival in late-stage compared with early-stage disease.^1,2^ While immunotherapy has transformed treatment paradigms in many cancers, its success in SCLC has been limited due to the unique challenges posed by the tumor’s immune microenvironment. PD-L1 blockade plus chemotherapy has become standard of care in late-stage SCLC and demonstrated prolonged overall survival, albeit only modestly.^3–9^

Unlike non-small cell lung cancer (NSCLC), SCLC has been considered an immune-cold tumor, typically associated with a lack of immune recognition, which promotes tumor progression and hinders effective immune responses.^1,2,10,11^ Recently, this paradigm of SCLC as an immune-cold tumor was challenged by its classification into 4 subtypes based on the expression of key transcription regulators. While SCLC subtypes of neuroendocrine origin, defined by expression of *ASCL1* (SCLC-A) and *NEUROD1* (SCLC-N), are generally considered immune-cold subtypes, a major non-neuroendocrine *POUF2F3-*driven subtype (SCLC-P) displays some inflammatory features. In addition, a more immune-infiltrated subtype (SCLC-I), typically of either neuroendocrine (NE) or non-neuroendocrine (non-NE) origin, has also been described.^1,8–10,12–14^

SCLC’s immunosuppressive tumor microenvironment (TME) is characterized by tumor-associated macrophages (TAMs) with a M2-like phenotype and is frequently infiltrated by regulatory T cells (CD4+ T_reg_ cells), collectively promoting tumor growth and contributing to immune evasion by establishing a suppressive network that limits immune-mediated tumor control.^6,15–23^ Importantly, the ratio of effector T cells to TAMs has recently been shown to impact the outcome of immune checkpoint blockade, with NE tumors with high T-cell and low macrophage abundance benefitting the most from anti-PD-L1 treatment.^20^

One major limitation of existing studies is the restricted access to patient material due to the aggressive nature of SCLC.^1,2,10,24^ While recent studies, including spatial analyses, have provided insights into the immune landscape and heterogeneity of SCLC, critical gaps remain in our understanding of immune cell diversity, function, and T cell recognition.^21^ The current lack of single-cell resolution and paired single-cell TCR sequencing (scTCR-seq) data represents a major gap knowledge of the multifaceted SCLC TME. A comprehensive view is critical for overcoming immunosuppressive barriers that restrain tumor-reactive T cells, ultimately improving the efficacy of immunotherapy in SCLC.^4–7^

Even though tumor-reactive T cell signatures have been successfully identified for many cancer types, including melanoma^25–27^, breast cancer^25^, colorectal cancer^25^, NSCLC^28^ and pancreatic adenocarcinoma (PDAC)^29^, a SCLC-specific signature is lacking, highlighting an urgent need to bridge this gap in the immunotherapeutic landscape. Moreover, linking those findings to SCLC subtypes is of utmost clinical relevance, since the modest survival benefit observed following checkpoint blockade plus chemotherapy appears to be subtype-dependent and most pronounced in the SCLC-I subtype.^8,9^ Improved understanding of immune engagement and restraint in SCLC is needed to enable hypothesis-driven combination strategies (e.g., checkpoint blockade and adoptive therapies), potentially complemented by approaches that enhance antigen presentation.^30^

Thus, we generated a high-resolution single-cell immune atlas of SCLC with paired αβ TCR sequencing, providing the most detailed view of the SCLC immune microenvironment to date. Linking transcriptional states with T cell clonality enabled the identification and validation of tumor-reactive TCRs in SCLC, marking the first direct evidence of T cell-mediated tumor recognition in this malignancy. Importantly, our study introduces a novel tumor-reactive T cell signature specific to SCLC. This signature not only differentiates reactive from non-reactive T cells but also correlates with patient survival and highlights key immune pathways, including *CXCL13* and *TNFRSF9* as markers of reactive T cells and *CTLA4* and *TIGIT* as additional immune checkpoints targets, laying the groundwork for next-generation immunotherapies in SCLC.

## Results

### Pan-lung cancer single-cell immune atlas reveals high patient variability and immunosuppression in SCLC

To characterize the immune landscape of SCLC tumors, we conducted paired single-cell RNA sequencing (scRNA-seq) and scTCR-seq on fluorescence-activated cell sorting (FACS)-purified CD45+ fractions from lung cancer patient biopsies (16 tumor samples from 14 SCLC patients, including tumors from primary lung tissues or lymph node metastases) (Fig. 1a, Suppl. Table S1). For comparative purposes, 3 primary tumor samples from NSCLC patients, and from large cell neuroendocrine carcinoma (LCNEC) patients, respectively, were also included (Fig. 1a, Suppl. Table 1). In total, the pan-lung cancer dataset is comprised of 61,201 single cells and 23,019 paired αβ-TCR chains. Unsupervised clustering of the 22 integrated samples created an immune landscape composed of 16 transcriptionally distinct clusters (Fig. 1b, Suppl. Fig. 1a), annotated based on canonical markers previously reported in literature (Suppl. Fig. 1b).

**Figure 1:**
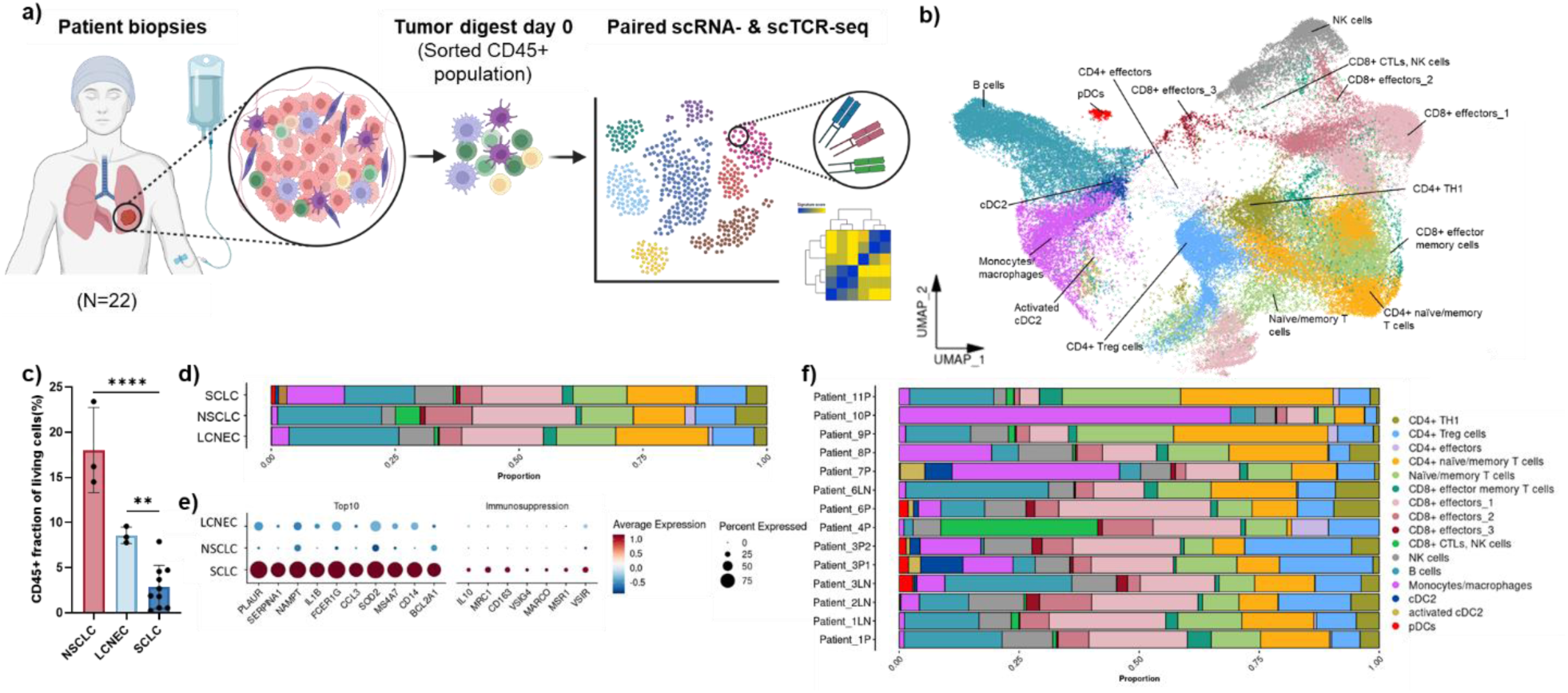
Single-cell immune profiling reveals high patient variability and immunosuppression in small cell lung cancer (SCLC) (a) Graphical summary of the experimental design for immune cell profiling (CD45+ cells) from tumour biopsies, encompassing 16 samples from 14 SCLC patients, 3 samples from 3 NSCLC patients, and 3 samples from 3 LNEC patients. Image created with BioRender.com. (b) UMAP of expression profiles of 61,801 single immune cells from tumour biopsies of patients mentioned in (a). Each dot on the UMAP represents a single cell and 16 broad immune cell subsets are annotated and marked by colour code. (c) The bar plot illustrates the frequency of CD45+ cells in tumour biopsies analysed by flow cytometry, with the CD45+ cell fraction represented on the y-axis as a proportion of all viable cells. (d) Comparison of cell-subset composition of immune infiltrates in tumour biopsies from pooled SCLC(n =14), NSCLC(n =3), and LCNEC (n =3) patients (same legend as in (f)). (e) Dot plot represents the top 10 differentially expressed genes between Monocytes/macrophages from SCLC versus NSCLC and LCNEC (p_val_adj < 0.05 & log2FC > 0.5, left side) and selected immunosuppressive markers for the same groups (right side). Color intensity represents scaled average gene expression, and dot size indicates the percentage of cells expressing each gene. (f) Immune cell-subset composition for each SCLC patient sequenced directly from the biopsies from primary tumours and lymph node metastases. For patient 3, two primary tumours and one lymph node metastasis were sequenced separately. (P – Primary tumour, LN – Lymph node metastasis)

A comparative analysis of SCLC, NSCLC, and LCNEC samples revealed a significantly lower immune cell compartment in SCLC patients compared with both NSCLC and LCNEC patients (Fig. 1c). A comparison of immune composition among different lung cancer entities showed that, while the monocyte/macrophage compartment was enriched in SCLC, CD8+ effector T cells and NK cell subsets were most abundant in NSCLC (Fig. 1d). Given the observed differences in immune cell composition, we next performed differential gene expression analysis on monocyte/macrophage populations in SCLC and compared them to those in NSCLC and LCNEC (Fig. 1e). Interestingly, we identified several immunosuppressive markers, including *PLAUR*, *SERPINA1*, *NAMPT*, *IL1B*, and *FCER1G*, among the top 10 upregulated genes in the monocyte/macrophage populations of SCLC patients with classical monocyte being the dominant myeloid subset (Fig. 1e, Suppl. Fig. 1c and d). These genes play pivotal roles in fostering a pro-tumorigenic immunosuppressive milieu, reinforcing mechanisms that may contribute to immune suppression in SCLC, as observed in previous studies.^31–35^

A closer investigation of immune cell distribution within SCLC patients revealed substantial intra-patient and inter-patient heterogeneity (Fig. 1f, Suppl. Fig. 2a-c). Patients 1, 3 and 6 provided both primary tumor (P) and lymph node metastasis (LN) samples, allowing for detailed intra-patient, site-specific analysis at single-cell resolution. In addition, for patient 3, two primary tumor regions (P1, P2) and a lymph node metastasis (LN) were analyzed. In patient 3, primary regions 1 and 2 exhibited a fivefold increase in monocytes/macrophages and a threefold increase in CD4+ T_reg_ cells compared to the lymph node (Suppl. Fig. 2b). Similarly, in patient 6, the primary tumor region demonstrated a fivefold increase in monocytes/macrophages, along with a two-fold increase in both CD8+ effector_1 and CD8+ effector_3 cells, and a threefold elevation in NK cells compared to lymph node metastasis (Suppl. Fig. 2c). Notably, the primary region was characterized by lower levels of CD8+ effector memory T cells, CD4+ naive/memory T cells, and B cells. The combined proportion of CD4+ T_reg_ cells and myeloid cells was consistently higher than that of T cells in the primary regions (Suppl. Fig. 2a-c). In line with the immunosuppressive phenotype of monocytes/macrophages in the SCLC TME (Fig. 1e, Suppl. Fig. 1d), these findings indicate a prevailing immune-cold, poorly infiltrated and tumor-promoting microenvironment across SCLC patients.

### Deep characterization of the T cell infiltrate unveils expanded CD8+ T cell subset with cytotoxic and exhausted features

Previous studies in various tumor types, including melanoma, renal cell carcinoma, and colorectal cancer have emphasized the importance of CD8+ effector T cells in orchestrating anti-tumor immunity, underscoring their therapeutic potential in enhancing immune-mediated tumor control.^36–42^ Importantly, recent work has explored single-cell T cell transcriptomes to pinpoint tumor-reactive T cells.^25–29^ For a deeper understanding of the SCLC-infiltrating CD8+ T cell biology and to identify tumor-reactive candidates and delineate their phenotype, we investigated their transcriptional states and TCR clonality.

An in-depth analysis of the transcriptional states of CD8+ effectors revealed four different transcriptional profiles in the SCLC TME (Fig. 2a). The first effector subpopulation, referred to as CD8+ effectors_1, displayed markers typically associated with cytotoxic CD8+ T cells, including *GNLY, NKG7, CD69, and KLRG1*, which have been described before in the context of CD8+ T-cell cytotoxicity and activation (Fig. 2b and c).^36,43–45^ The second subpopulation, which we designated as CD8+ effectors_2, was defined by the expression of both cytotoxic and exhaustion markers, with a cytotoxic gene signature similar to that of CD8+ effectors_1 but enriched for exhaustion-associated markers such as *HAVCR2*, *ENTPD1*, *TOX*, *TIGIT*, and *CTLA4* (Fig. 2b and c).^44^ Such a co-expression of cytotoxic and exhaustion markers has previously been linked to tumor-reactivity.^25–29^ The third subpopulation, which we termed CD8+ effectors_3, was marked predominantly by the expression of proliferation-associated genes, including *TOP2A*, *BIRC5*, *CDC20*, *TK1*, and *MKI67*, suggesting a cycling subset of CD8+ T cells rather than a distinct differentiation state (Fig. 2b and c). Lastly, a forth population exhibited expression of markers typically associated with a memory- or stem-like phenotype, such as *TCF7, CCR7, LEF1* and *IL7R,* and was accordingly termed CD8+ effector memory T cells (Fig. 2b and c).^44,45^

**Figure 2:**
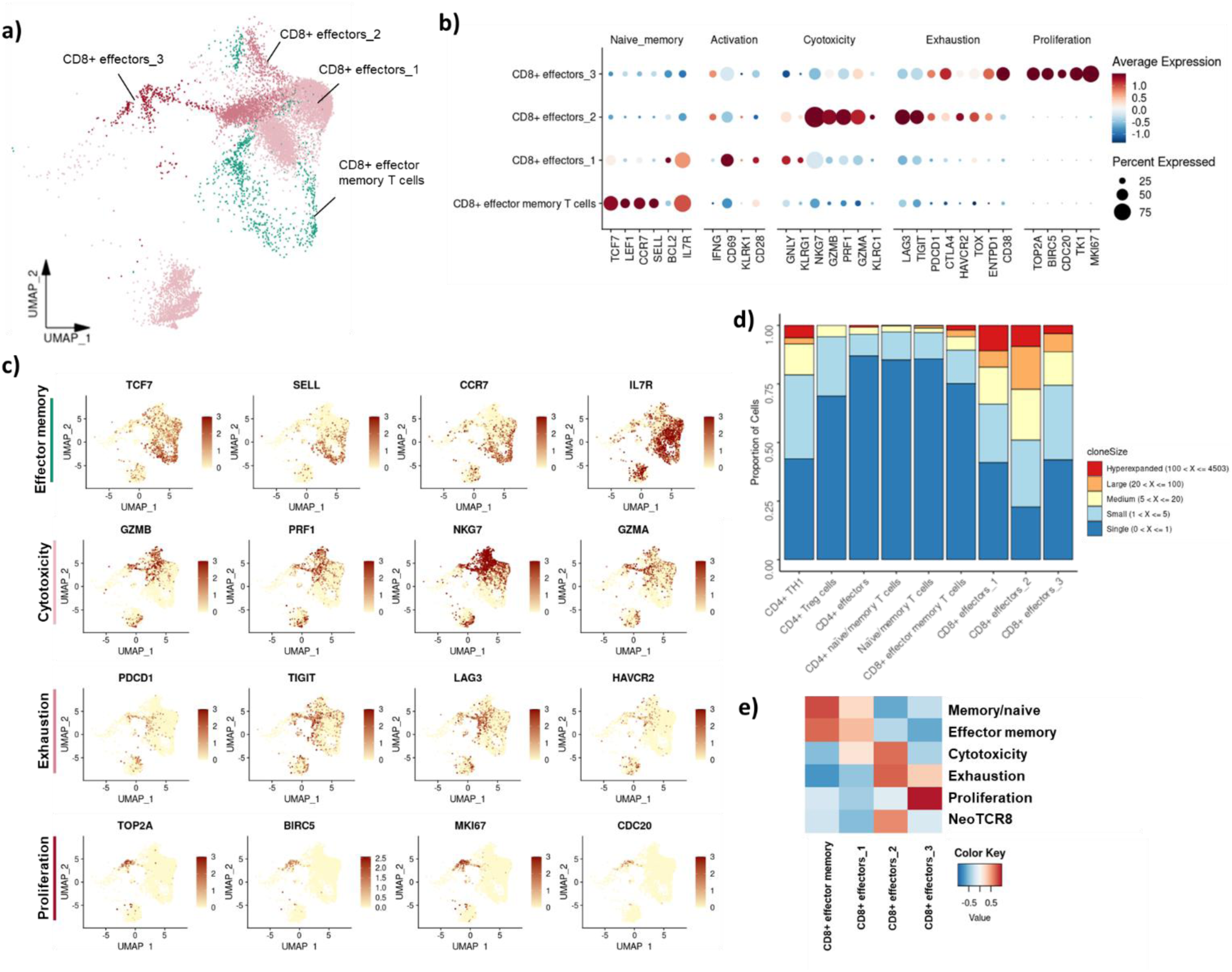
Combined scTCR-seq and scRNA-seq analysis reveals three transcriptional CD8+ effector states. (a) CD8+ T cells from SCLC biopsies are subsetted from the main UMAP shown in Fig. 1b. (b-c) Cytotoxic, exhaustion and proliferation gene markers expressed by different CD8+ effector states are depicted in a dotplot (b) and a feature plots (c). (d) The bar plot represents the distribution and clonality of TCRs on T cells from the SCLC tumor microenvironment. Clonality is quantified as clone size across different T cell states. Clone size refers to the number of cells expressing a specific TCR. (e) The heat map visualization shows the gene-set variation analysis of CD8+ TILs from SCLC patients. The y-axis displays the applied gene signatures, while the x-axis represents the selected CD8+ T cell subsets within the SCLC tumor microenvironment. In addition to known T cell phenotypic gene panels, the NeoTCR8 signature from Lowery et al. (Science, 2022) was applied to the CD8+ TILs.

We sequenced 15,627paired αβ-TCRs from all 14 SCLC patients and investigated the cell subset-specific distribution and clonality of TCRs (Fig. 2d). While clonotypes predominantly originating from memory or naïve T cell states, including naïve/memory T cells, CD4+ naïve/memory T cells, and CD8+ effector memory T cells showed little to no clonal expansion, the CD8+ effector T cell compartment and, to some extent, CD4+ TH1 cells, were enriched for expanded TCR clonotypes. Notably, CD8+ effectors_2, followed by CD8+ effectors_1, represented the most clonally expanded subsets (Fig. 2d). These findings were further supported by an intra-patient analysis across multiple sites for 3 patients, which showed that although CD8+ effectors_2 harboured the highest proportion of clonally expanded T cells within the shared clones in the majority of cases (Suppl. Fig. 2d, upper panel), the CD8+ effectors_1 subset contained the highest number of expanded TCR clonotypes (Suppl. Fig. 2d, lower panel).

Bystander T cells can also undergo expansion due to unrelated antigenic stimuli, prompting the need to distinguish true tumor-reactive T cells from bystanders.^25^ We therefore applied the previously described NeoTCR8 signature, reported to predict tumor-reactivity, which was derived from 14 metastatic samples in melanoma, breast, and colorectal cancers, encompassing 46 tumor-reactive and 3 non-tumor-reactive T-cell clones.^25^ Strikingly, applying this signature to our dataset revealed significant enrichment of NeoTCR8 signature genes within the CD8+ effectors_2 population, suggesting that this subset may represent a key reservoir of tumor-reactive T cells (Fig. 2e).

Overall, the in-depth characterization of CD8+ T cell states and TCR clonotypes identified CD8+ effector_2 cells as principal subset containing tumor-reactive T cell candidates by showing co-expression of cytotoxic and exhaustion markers, one of the highest clonality as well as highest enrichment in a publicly available tumor-reactive signature.

### Tumor-reactive TCRs were functionally validated through patient-derived cell line recognition

Having highlighted CD8+ effectors_2 cells as the candidate subset potentially harboring a high number of tumor-reactive TCRs, we next functionally validated tumor-reactive T cells using autologous cell lines generated from patient-derived xenograft (PDX) models from tumor biopsies of SCLC patients 5, 6 and 7 (Suppl. Fig. 3a).

Whole exome sequencing was employed to delineate the mutational landscape, demonstrating a high somatic mutation burden in the parental tumor as well as cell lines (Suppl. Fig. 3b). Furthermore, over 70% of the mutations were shared between the parental tumor and the PDX-derived spheroid cell lines, indicating that these cell lines retained the majority of patient-specific mutations and are suitable for tumor-reactivity testing (Suppl. Fig. 3c). We furthermore sought to determine on RNA expression level, whether the cell lines matched any of the established SCLC subtypes.^1,2,9,10^ The cell lines derived from patients 6 and 7 exhibited high *ASCL1* expression, consistent with of the SCLC-A subtype, whereas the cell line from patient 5 displayed a mixed expression profile of *ASCL1* and *NEUROD1*, more consistent with the SCLC-N subset. These tumor subsets are typically associated with immune-cold phenotype and low immune infiltration (Suppl. Fig. 3d).

First, to assess the effect of tumor cell–mediated stimulation on the T cell subset distribution and T cell reactivity, we expanded tumor-infiltrating lymphocytes (TILs) in the presence of autologous tumor cells from patient 6 (Suppl. Fig. 4a) and we observed that CD8+ effectors_2 exhibited the most efficient expansion compared to culture conditions without tumor cell stimuli (Suppl. Fig. 4b and c). Moreover, co-culture with autologous tumor cells resulted in the highest tumor-killing capacity (Suppl. Fig. 4d), further corroborating experimentally the transcriptional evidence (Fig. 2b-e) that the CD8+ effectors_2 cells might contain the main tumor-responsive CD8+ T cell population.

Next, given the large number of candidate TCRs within the CD8+ effectors_2 subset, we implemented a ranking strategy prior to functional *in vitro* validation to refine our selection of tumor-reactive clonotypes by ranking TCRs from this population in 3 patients based on NeoTCR8 signature scores (Fig. 3a). During this process, virus-specific TCRs were identified using publicly available datasets^46^ and excluded to avoid bias from bystander T cell responses (Suppl. Fig. 5a). After removing predicted virus-specific TCRs, we tested a set of 150 TCR candidates (top 50 ranked per patient) that were strongly enriched for tumor-reactive features.

**Figure 3:**
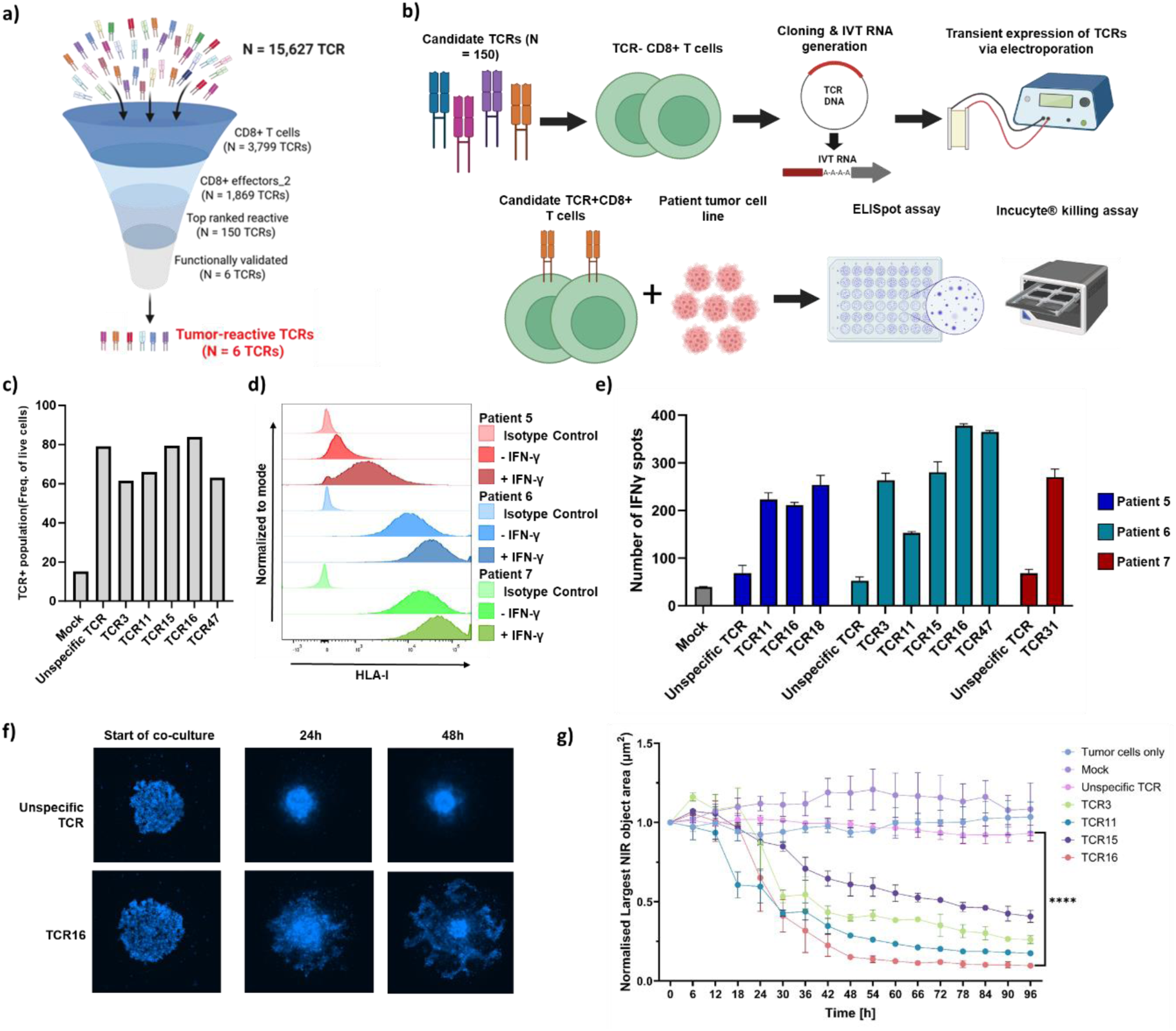
Identification, prioritization, and functional validation of tumor-reactive TCR repertoires from SCLC patients. (a) The schematic representation summarizes the stepwise prioritization and validation process, narrowing down from an initial pool of 15,627 TCRs to 6 functionally validated tumor-reactive TCRs. (b) Experimental setup for the functional validation of TCR candidates, shortlisted using the NeoTCR8 signature, is depicted. Endogenous TCRs from CD8+ T cells from healthy donors are disrupted using CRISPR followed by transfection with mRNA encoding for TCR candidates. Upon co-culture of T cells and autologous cell lines T cell reactivity was measured in ELISpot and killing assays. Image created with BioRender.com. (c) Flow cytometry data illustrates the expression of candidate TCRs on CD8+ T cells from healthy donors. (d) Flow cytometry data shows MHCI expression on patient-derived xenografts (PDX) from SCLC patient 5, 6 and 7 before and after IFN-γ treatment. (e) IFN-γ ELISPOT assay results demonstrating the immune response elicited by CD8+ T cells from healthy donors transfected with our shortlisted TCR candidates from Patients 5, 6 and 7. (f) Images from the 3D tumor spheroid killing assay for TCR16 from patient 6, where tumor cells are stained in infrared dye (blue), demonstrate disintegration of tumor spheroids when tumor-reactive TCR-expressing T cells are introduced (lower panel), compared to control T cells with non-specific TCRs (upper panel). (g) Tumor spheroid integrity was followed over time in a 3D tumor-killing assay for five TCRs from patient 6. The y-axis shows normalized spheroid intensity (size of the largest object in each well), and the x-axis represents the assay duration. Different colors represent each TCR, as indicated in the legend on the right.

For functional validation, candidate TCRs were cloned into appropriate vector backbones for *in vitro* mRNA transcription. Subsequently, these TCRs were transiently expressed in CD8+ T cells from healthy donors, in which endogenous TCRs had been knocked out using CRISPR gene editing technology (Fig. 3b). Efficient disruption of the endogenous TCR with about 80% and expression of the targeted TCRs were confirmed prior to functional assays (Fig. 3c, Suppl. Fig. 5b). For functional validation, we used autologous tumor cell lines deriving from the corresponding patients (Fig. 1a, Fig. 3b). Given the baseline (patient 5) expression of human leukocyte antigen (HLA)-I, tumor cells from all 3 patients were pre-treated with interferon-gamma (IFN-γ) to enhance HLA-I upregulation before the assays (Fig. 3d). Tumor-reactivity was first assessed using an IFN-γ-based Enzyme-Linked Immunospot (ELISpot) assay by co-culturing TCR-transfected CD8+ T cells with autologous tumor cells. Of the 50 TCRs screened per patient 5, 6 and 7 (150 TCRs in total), 9 clones displayed functional reactivity, as evidenced by the number of IFN-γ spots (Fig. 3e, Suppl. Fig. 5c). To further validate tumor-specificity, we performed a second screening using transgenic Jurkat reporter cells and K562 as orthogonal antigen-presenting cells (APCs) transiently expressing patient-specific HLAs. This approach allowed us to identify 3 off-target TCRs, which were excluded from further analyses (Suppl. Fig. 5d). Finally, to obtain direct evidence of tumor cell lysis, we conducted a 3D tumor spheroid killing assay for patients 5 and 6 in a third screening. This assay demonstrated tumor spheroid disintegration upon co-culture, confirming tumor cell killing (Fig. 3f and g, Suppl. Fig. 5e and f). Overall, 6 of the 9 TCRs that were reactive in ELISpot assays and confirmed to be tumor-specific demonstrated strong functional potency, effectively killing tumor cells.

To follow tumor-reactive T cells within the different CD8+ T cell subsets, we annotated tumor-reactive TCRs within the combined scRNA-seq and scTCR-seq dataset based on our functional validation. In patient 6, we see the highest proportion and clonality of tumor-reactive T cells in the CD8+ effectors_2 state (Suppl. Fig. 5g), reinforcing the notion that this population might be critical for identifying a robust tumor-reactive T cell signature in SCLC but not restricted to this subset.

Notably, despite their high ranking according to the NeoTCR8 signature, the top 2 TCRs, TCR1 and TCR2, did not display detectable tumor reactivity in ELISpot assays. In contrast, TCR16, despite ranking lower, emerged as one of the most potent tumor-reactive TCRs (Fig. 3e). These findings suggest that while the signature enriched for tumor-reactive candidates, further refinement may improve its predictive performance in SCLC datasets.

### Transcriptional analysis of tumor-reactive CD8+ T cells reveals upregulation of distinct signature genes with biological implications in SCLC TME

To define a molecular signature in tumor-reactive T cells specifically for SCLC, we compared the differential gene expression between reactive and non-reactive T cells within CD8+ T cells from SCLC tumor biopsies (Fig. 4a). Here, we excluded untested CD8+ effectors_2 cells from the analysis to avoid potential discrepancies due to potentially tumor-reactive TCRs within this population. In a second refinement step, we filtered for signature genes that were exclusively upregulated in tumor-reactive T cell compared to non-reactive T cells giving rise to 43 genes, designated as ‘SCLC_TR’ (Fig. 4b). Application of the SCLC_TR signature to the SCLC dataset revealed an enrichment within a distinct subset of the CD8+ effectors_2 population (Fig. 4c). In contrast, non-reactive T cell signatures were predominantly expressed in CD8+ effector memory T cells and CD8+ effectors_1 subsets (Suppl. Fig. 6a). Leveraging our signature to predict tumor-reactive CD8+ T cells, including those that were not functionally validated, revealed that the predicted tumor-reactive repertoire predominantly resides within the CD8+ effectors_2 population, whereas most non-tumor reactive clones are enriched in the CD8+ effectors_1 population (Suppl. Fig. 6b). This pattern mirrors the distribution of viral TCRs (Suppl. Fig. 5a), suggesting that bystander T cells may predominantly reside within the CD8+ effectors_1 state, further distinguishing tumor-reactive from non-reactive T cell populations.

**Figure 4:**
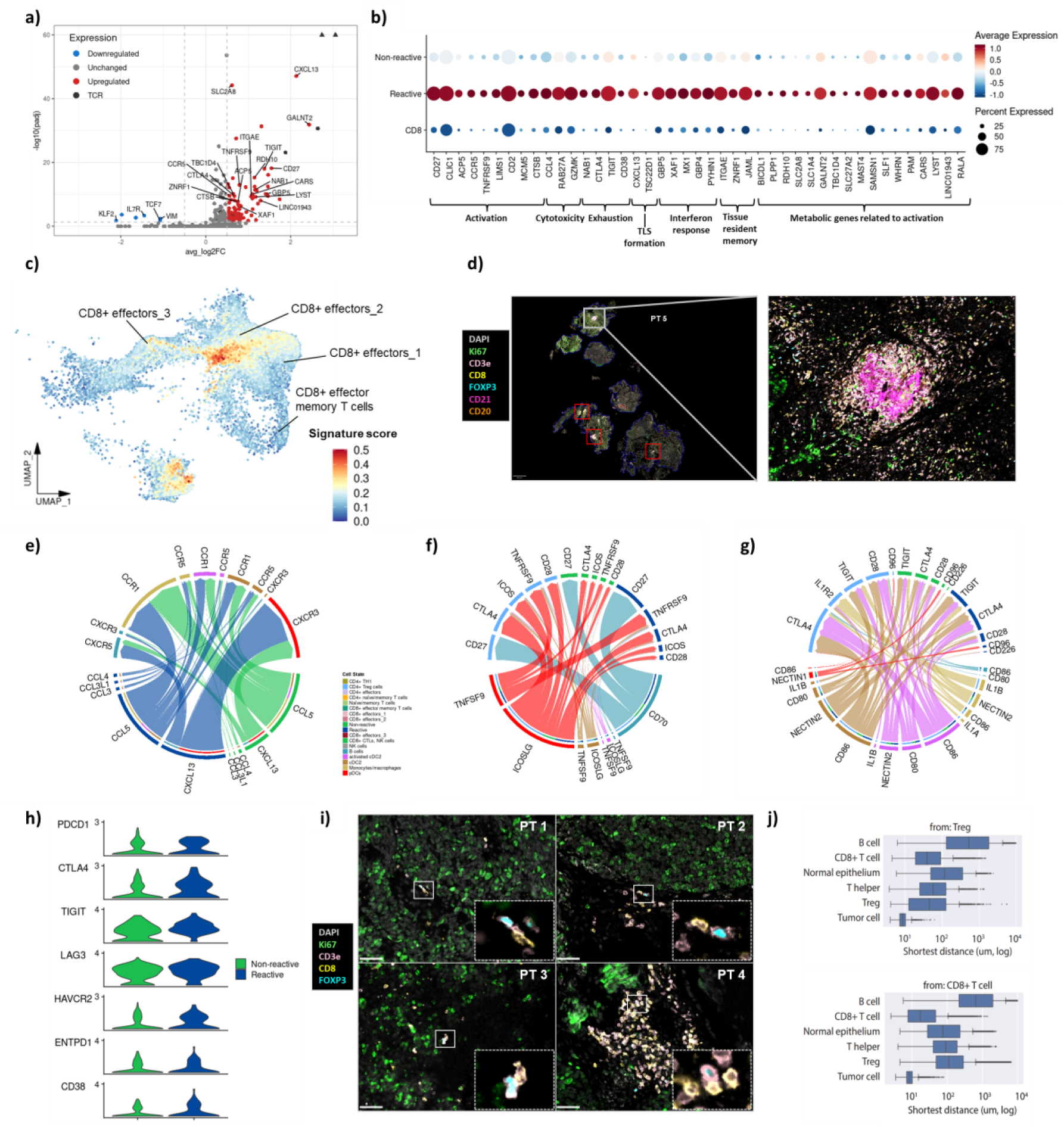
Unveiling a distinct molecular signature in tumor-reactive CD8+ T cells and characterizing its functional implication in SCLC TME. (a) The volcano plot shows genes differentially expressed between reactive T cells and non-reactive CD8+ T cells from SCLC biopsies. CD8+ effector_2 cluster was excluded from analysis to avoid bias from potential untested tumor-reactive TCRs. Upregulated genes in reactive T cells appear on the right side of the volcano plot, while downregulated genes are on the left. The legend on the bottom right depicts the colors used to represent gene regulation, and specifically in black, genes related to the TCR alpha and beta chains (filtering criteria for DGEA: p_val_adj < 0.05 & log2FC > 0.5). (b) The dot plot visualizes the expression of molecular signature genes from significantly upregulated genes in reactive T cells and removal of non-significant genes between reactive and non-reactive T cells (p_val_adj < 0.05 & log2FC > 0.35). Expression of signature genes is compared across reactive T cells, non-reactive T cells, and other CD8+ subsets (excluding CD8+ effectors_2). The color gradient represents the magnitude of average expression, ranging from low (blue) to high (red). The size of the circles reflects the fraction of cells expressing the gene in percentage. The genes are ordered according to function. (c) The feature plot shows the spatial distribution across the UMAP of subsetted CD8+ T cells from SCLC patient biopsies enriched in the SCLC_TR signature of tumor-reactive T cells. (d) Representative high-plex immunohistochemistry (IHC) staining illustrates the spatial distribution of mature TLS structures within a SCLC tumor sample with a magnified view of a representative TLS characterized by a core of B cells (CD20+) and follicular dendritic cells (CD21+) surrounded by a ring of CD4+ T_reg_ cells (CD3+ FOXP3+ CD8-), T helper cells (CD3+ FOXP3− CD8−) and cytotoxic CD8+ T cells (CD3+ CD8+) in a peritumoral region (Ki67+) (scale bar: 1.5 mm; inset scalebar: 50 µm). (e) Cell-cell communication analysis showcases elevated predicted chemokine and cytokine signaling between reactive and non-reactive T cells on the bottom as source cells and pDCs, monocytes/macrophages, activated cDC2, cDC2, and B cells on the top as target cells in all SCLC patients. (f-g) Cell-cell communication analysis showcases elevated predicted immunomodulatory ligand-receptor interactions between CD4+ T_reg_ cells, reactive and non-reactive T cells as target cells and pDCs, monocytes/macrophages, activated cDC2, cDC2, and B cells as source cells in all SCLC patients. The circos plot depict either co-stimulatory signaling (f) or co-inhibitory signaling (g), respectively (same legend as in (e)). (h) Violin plots depict the expression of inhibitory receptor expression across tumor-reactive and non-reactive CD8+ T cell subsets. (i) Representative high-plex IHC staining shows the spatial proximity of CD4+ T_reg_ cells (CD3+ FOXP3+ CD8-) and CD8+ T cells (CD3+ CD8+) within SCLC tumor sections from four different patients (scale bar: 50 µm). (j) Faceted box plots illustrate the distribution of average shortest distances (µm) from each cell phenotype to target cell phenotype CD4+ T_reg_ cells (upper panel) and CD8+ T cells (lower panel), respectively.

Even though we confirmed the presence of tumor-reactive T cells within SCLC tumors, clinical responses to immunotherapy remain modest. This suggests that additional constraints might limit effective T cell activity, prompting us to examine the contributions of individual signature genes in greater detail. The SCLC_TR signature comprises genes associated with T-cell effector function (e.g., CTSB, GZMK), T-cell exhaustion (e.g., CTLA4, TIGIT), tertiary lymphoid structure (TLS) formation (e.g., CXCL13), T-cell activation (e.g., CD27, TNFRSF9), and tissue residency (e.g., ITGAE) (Fig. 4b).^25–29^ In addition to these markers, the signature includes several metabolism-related genes, such as SLC1A4, SLC2A8, SLC27A2, TBC1D4, RALA, and PLPP1, which are linked to amino acid uptake, glucose transport, lipid handling, and mTORC1 signalling—pathways essential for sustained CD8⁺ T-cell activation.^47–50^ Furthermore, genes involved in retinoid metabolism (RDH10) and lysosomal or trafficking pathways (LYST, BICDL1) suggest that tumor-reactive T cells in SCLC undergo coordinated metabolic rewiring to support anabolic growth, effector function, and adaptation to the metabolically restrictive TME.^47,51^ Notably, *CXCL13*, a key player in TLS formation and B cell and T cell communication, has been identified as a marker for tumor-reactive T cells in clinical trials.^52^ Its expression has been associated with improved prognosis and enhanced survival outcomes across multiple cancer types, further underscoring its significance in shaping effective anti-tumor immune responses.^52–55^ Further, *CXCL13* is commonly associated with tumor-reactivity in other solid cancers.^25,27–29,56^ Consistent with those previous reports, we could observe the formation of multiple mature TLS in the TME of a single patient in an independent cohort of SCLC patients (Fig. 4d, Suppl. Fig. 6c) underlining the relevance of *CXCL13* and mature TLS for tumor-reactive CD8+ T cells also in SCLC. We then performed cell-cell interaction analysis to analyze cytokine and chemokine-receptor interactions within the tumors. By defining reactive and non-reactive T cells as senders and antigen-presenting cells (APCs) as receivers in the TME, *CXCL13–CXCR5/CXCR3* signaling emerged as the most prominently predicted interaction in reactive T cells compared with non-reactive T cells, mediating interactions specifically with B cells and dendritic cells (Fig. 4e). In contrast, the *CCL3/CCL4-CCR5* axis was the most prominent interaction when tumor-reactive T cells were set as receivers (Suppl. Fig. 6d). *CCR5* is also one of the novel markers in the SCLC_TR signature, next to *MX1, GZMK, GBP5, CD38*, which are associated with T cell activation and exhaustion and are not included in any previously defined molecular tumor-reactive T cell signature (Fig. 4b).^57^ The *CCL3/4–CCR5* axis was reported to facilitate CD8+ T-cell infiltration and activation in tumors, enhancing cytotoxic responses, but persistent signaling may contribute to immune exhaustion.^57–59^

To further elucidate the intercellular communication of reactive and non-reactive T cells within the SCLC TME, we analyzed co-stimulatory and co-inhibitory ligand-receptor interactions (Fig. 4f and g). In this context, CD4+ T_reg_ cells were included given their known immunosuppressive role in tumor immunity.^60^ APCs, namely B cells and myeloid cells, were once more chosen as key signaling counterparts and dominant co-stimulatory ligand-receptor interactions (*TNFSF9-TNFRSF9* and *CD70-CD27*) were identified, with more pronounced predicted interactions in reactive T cells compared to non-reactive ones (Fig. 4f). The *TNFSF9–TNFRSF9* interaction, originating from myeloid cells, is reported to enhance the CD8+ T cell response by activating *TNFRSF9* on their surface.^61^ Similarly, the *CD70–CD27* axis, driven by B cells, is essential for CD8+ T cell activation, survival, and memory formation.^62^

Furthermore, several dominant co-inhibitory ligand-receptor interactions (*NECTIN2-TIGIT* and *CD86-CTLA4*) were identified with increased interaction probabilities in reactive T cells compared to non-reactive T cells (Fig. 4g). The *NECTIN2–TIGIT* axis functions as an immune checkpoint, dampening T-cell activation via *TIGIT*, which is upregulated in exhausted T cells, thereby limiting cytotoxic function.^63–65^ We further examined immune checkpoint expression and, in addition to *CTLA4, TIGIT* and *CD38,* we identified *HAVCR2, LAG3, PDCD1, and ENTPD1*, all of which were expressed at higher levels in reactive T cells compared with non-reactive T cells within the SCLC TME (Fig. 4h). Interestingly, *CD38* has recently been described as novel immune checkpoint linked to exhaustion and dysfunction in CD8+ T cells and decreased mitochondrial fitness.^66–68^

In addition, several co-stimulatory and co-inhibitory interactions (*TNFSF9-TNFRSF9, ICOSLG-ICOS, IL1B-IL1R2*, *CD80/CD86–CD28/CTLA4 and NECTIN2-TIGIT* axes) between myeloid cells and CD4+ T_reg_ cells were predicted (Fig. 4f and g). This immunomodulation of CD4+ T_reg_ cells suggests an enhancement of their suppressive function and limiting effector T cell activation, a well-established immune checkpoint mechanism in cancer.^61,63–65,69–71^ In line with this predicted suppressive interaction, we could observe colocalization between T_reg_ and CD8+ T cells in the TME across all four patient samples of an independent cohort using multiplex immunofluorescence (Fig. 4i and j). This close proximity underscores the potential crucial role of CD4+ T_reg_ cells to suppress T cell activity within the tumor.

Lastly, the proportion of predicted tumor-reactive CD8+ T cells tended to increase with the abundance of myeloid and CD4+ T_reg_ cells, as observed for patient 3 and 7 (Suppl. Fig. 6e and f). Taken together with our comparative analysis of SCLC, NSCLC, and LCNEC TMEs (Fig. 1e), which demonstrated that monocytes/macrophages in the SCLC TME exhibit strong immunosuppressive properties, these findings reinforce the notion of an immunosuppressive mechanism counteracting tumor-reactive T cell activation, potentially dampening their anti-tumor response.

Collectively, these findings highlight the complex interplay between tumor-reactive T cells, myeloid compartments, and CD4+ T_reg_ cells, that seem to counterbalance the activation of antitumor T cells via direct and indirect immunosuppression in the SCLC microenvironment in a synergistic manner.

### SCLC_TR identifies successfully and robustly tumor-reactive TCRs in SCLC and PDAC

Next, we evaluated the success rate of our SCLC_TR signature and explored the overall performance against other signatures. First, we benchmarked the gene signature performance *in vitro* by reranking TCR sequences according to their potential tumor-reactivity using the SCLC_TR signature and we identified 47 additional tumor-reactive TCR candidates using our IFN-γ-based ELISpot by co-culturing TCR-transfected CD8+ T cells with autologous tumor cells from 2 patients (Fig. 5a). Strikingly, SCLC_TR outperformed NeoTCR8 in identifying tumor-reactive TCRs, achieving a success rate of approximately 32% versus 6% (Fig. 5a). In addition, we observed a high proportion of tumor-reactive TCRs in metastatic sites (lymph node metastasis for patient 6 and brain metastasis for patient 7), whereas the proportion remained comparably low at the primary tumor site (patient 6) (Fig. 5b).

**Figure 5:**
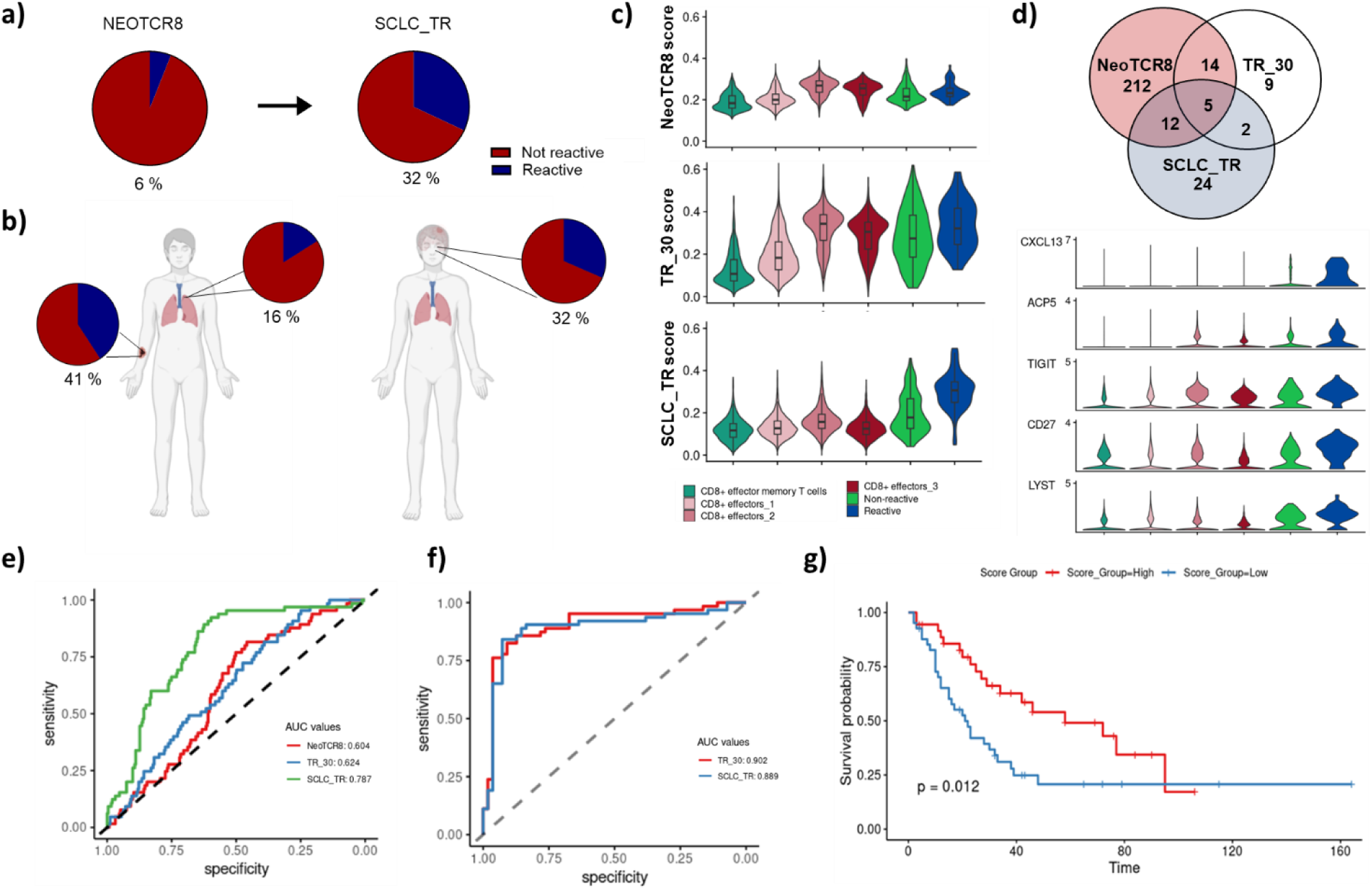
Assessment of SCLC_TR success rate and benchmarking its performance in SCLC and PDAC. (a) Pie charts next to patients depict the percentage of tumor-reactive and non-reactive TCRs evaluated using two patient-derived cell lines. Overall success rates for the identification of tumor-reactive TCRs in patient 6 and 7 are shown based on either the scoring using the NeoTCR8 or the SCLC_TR gene signature, respectively. Image created with BioRender.com. (b) Schematic representation of tumor-reactive proportions for SCLC_TR-ranked and validated TCR candidates per tumor sites. For patient 6, TCR sequences derived from the primary site (lung) and lymph node metastasis were evaluated whereas for patient P7 only TCRs derived from a brain metastasis were tested by IFN-γ-based ELISpot. (c) Violin plots illustrate the expression of three distinct gene signatures in CD8⁺ TILs from SCLC patients, with the top, middle, and bottom panels corresponding to the NeoTCR8, TR_30, and SCLC_TR signatures, respectively. (d) The Venn diagram shows the overlap of number of genes between, TR_30 and SCLC_TR (upper panel). Violin plots show the expression of the 5 genes overlapping across all the molecular signatures (NeoTCR8, TR_30 and SCLC_TR, lower panel, same legend as in (c)). (e) The ROC analysis evaluates the performance of NeoTCR8 and TR_30, from Lowery et al. and Meng et al., respectively, to our delineated SCLC_TR signature on our SCLC dataset. (f) The ROC analysis evaluates the performance of different signatures from the literature and our signature on external PDAC tumor dataset. (g) Performance evaluation of SCLC_TR on bulk RNA-seq data from George et al. Red survival curve represents patients which are high for SCLC_TR whereas the survival curve in blue represents patients which were scored to be low for the SCLC_TR.

To evaluate the efficacy of the tumor-reactive signature SCLC_TR beyond our dataset, we first compared its performance with 2 external signatures from recent studies that aimed to delineate tumor-reactive signatures across different tumor types.^25,29^ The NeoTCR8 signature derived from metastatic cancer samples including melanoma, breast, and colorectal cancers was derived from tumors commonly considered as infiltrated “hot” tumors.^25^ In contrast, the TR_30 signature, comprising 93 tumor-reactive and 65 non-tumor-reactive T cell clones, was derived from 9 PDAC cases, generally classified as immune-cold tumors.^29^ A comparison of the enrichment of our SCLC_TR signature with the NeoTCR8 and TR_30 signatures in CD8+ T cells, including both reactive and non-reactive T cells, revealed that all signatures were most strongly enriched in the CD8+ T cell effectors_2 subset (Fig. 5c). However, SCLC_TR showed the clearest separation between reactive and non-reactive T cells (Fig. 5c).

Several SCLC_TR signature genes overlap with previously reported tumor-reactive signatures, with 17 genes shared with NeoTCR8, 7 genes shared with TR_30 (Fig. 5d, Suppl. Fig. 7a and b). 5 genes, CXCL13, *ACP5, TIGIT, CD27,* and *LYST*, were consistently shared across all 3 signatures (Fig. 5d). Among them, *ACP5* and especially, *CXCL13* showed almost exclusive upregulation in reactive T cells, underscoring their potential as markers for tumor-reactivity (Fig. 5d). While *ACP5* remains functionally poorly defined, CXCL13 has a well-established role in anti-tumor immunity and is consistently associated with tumor-reactive T cells across multiple solid cancers^25,27–29,52^

Importantly, this superior performance of SCLC_TR in identifying tumor-reactive CD8+ T cells compared with the other 2 signatures in our SCLC dataset was confirmed by ROC curve analysis (Fig. 5e). To further assess the performance of SCLC_TR, we applied it to the Meng et al. PDAC dataset, from which the TR_30 signature was originally derived from.^29^ Remarkably, SCLC_TR performed comparably to TR_30, indicating that it can robustly identify tumor-reactive CD8+ T cells across datasets from other cold TMEs (Fig. 5f). We further evaluated the performance of SCLC_TR using bulk RNA sequencing data from a clinical SCLC cohort reported by George et al.^2^ (Fig. 5g). Strikingly, patients with higher SCLC_TR signature scores exhibited significantly improved survival compared with those with lower scores, suggesting potential prognostic relevance.

In summary, the SCLC_TR signature robustly identifies the tumor-reactive CD8+ T cell repertoire in SCLC with remarkable reliability and may represent a promising new biomarker of patient survival. Moreover, the presence of tumor-reactive CD8+ T cells may contribute to a sustained anti-tumor immune response, potentially enhancing survival outcomes in these patients.

## Discussion

This study provides an extensive single-cell analysis of the immune landscape in SCLC, deeply characterizing a so far still considered immune-cold TME, revealing marked immunosuppression and significant heterogeneity both within and between patients. By integrating scRNA-seq and scTCR-seq, we uncovered tumor-reactive T cells that effectively recognized and eradicated tumor cells. We provide a SCLC_TR signature to identify such tumor-reactive T cells with high fidelity.

Previous studies have characterized SCLC as an immune-cold tumor with limited immune infiltration, particularly the SCLC-A and SCLC-N subtypes, to which the patients in our study predominantly belong to (Suppl. Fig. 3c).^1,2,10^ Consistent with this classification, our analysis demonstrated a significantly reduced immune cell compartment in SCLC compared to NSCLC and LCNEC, including lower proportions of NK cells and cytotoxic CD8+ T-cell subsets alongside an enrichment of monocyte/macrophage populations.^18,19,21,22,72^ Differential gene expression analysis further highlighted the upregulation of *PLAUR, SERPINA1, IL1B, NAMPT,* and *FCER1G* in SCLC-associated monocyte/macrophage populations, described as immunosuppressive-associated markers, described to contribute to immune evasion, T_reg_ cell survival and tumor progression.^31–35^ *PLAUR* facilitates extracellular matrix degradation and immune evasion, while *SERPINA1* inhibits pro-inflammatory proteases, enhancing anti-inflammatory responses and regulatory T cell survival.^32,33^ *IL1B* drives chronic inflammation that recruits MDSCs and supports TAM-mediated immune suppression.^73–75^ *NAMPT* enhances metabolic fitness and anti-inflammatory cytokine production under hypoxic conditions, further sustaining immunosuppressive TAMs.^31^ *FCER1G* supports immune signalling pathways that suppress T-cell activation and contribute to tumor progression.^30^

The aggressiveness of SCLC and limited biopsy access complicate the collection of patient samples, hindering research on its immune landscape.^1,2,10^ Previous research has primarily focused on tumor heterogeneity and stromal interactions rather than providing an in-depth characterization of immune cells, particularly T cells, within the SCLC TME.^18–22,24^ Furthermore, studies have largely analyzed limited patient cohorts, restricting the generalizability of their findings and leaving critical questions about the functional diversity and exhaustion states of T cells unanswered.^18–22^ To our knowledge, our study hitherto provides one of the most extensive single-cell immune atlases of SCLC, coupled with paired TCR sequencing that enables in-depth T cell-focused analyses. The combined scRNA-seq and scTCR-seq data unveiled 3 different transcriptional states in CD8+ effector T cells, spanning early effector-like cytotoxic CD8+ effectors_1, effector-like exhausted CD8+ effectors_2 and proliferative CD8+ effectors_3. Our findings suggest that CD8+ effectors_2 represent a critical effector population, with concordant expression of cytotoxic and exhaustion signatures and the highest T cell clonality. This marks them as a key subset in anti-tumor immunity providing a framework to identify new potential biomarkers or therapeutic targets, warranting further investigation of their functional relevance. These observations are in line with previous reports showing that clonally expanded TCRs often harbor tumor specificity, underscoring their utility as biomarkers and therapeutic targets.^76^

From 15,627 TCRs, 150 were shortlisted for functionality studies using the NeoTCR8 signature^25^ as this was the most compelling signature to date when our study started. During the course of this work, the PDAC-derived TR_30 signature was published and provided an additional external benchmark.^29^ The NeoTCR8 signature was enriched in the CD8+ effectors_2 population, ultimately leading to the identification of 6 tumor-specific TCRs. Functional assays validated the tumor-killing capacity of these TCRs, with TCR-transfected CD8+ T cells exhibiting enhanced cytotoxicity and specificity when co-cultured with autologous tumor cells. This finding is the first description of tumor-reactive TCRs for SCLC challenging the notion of a so far immune-cold tumor as we show efficient tumor reactivity and clearance. These findings establish a foundation for future efforts to develop immunotherapeutic strategies tailored to SCLC.

Our screening method, unlike conventional approaches such as reporter-based assays, cytokine release assays, peptide-major histocompatibility complexes (pMHC) multimer staining, or bulk TCR sequencing, does not depend on patient-derived PBMCs. Instead, it utilized PBMCs from healthy donors and was conducted against patient-derived tumor cell lines, enabling standardized functional comparison across candidate TCRs while reducing confounding factors from patient-specific immune variability.^77^ This approach also offers significant advantages, particularly for patients or tumor types where obtaining blood samples may be challenging. By bypassing the dependency on patient-derived immune cells, this method ensures that robust screening for tumor-reactive TCRs can still be conducted. Additionally, this strategy avoids variability introduced by patient-specific immune dysfunction, which is often a hallmark of SCLC and other cold TMEs. The inclusion of functional co-culture assays to measure IFN-γ secretion (ELISpot), killing (live-cell imaging) and to exclude off-target TCRs (K562 and NFAT-Jurkat co-culture assay), allowed for the identification of TCRs with genuine anti-tumor efficacy and tumor-specificity, providing a higher level of stringency compared to traditional TCR screening methods.

To better characterize tumor-reactive CD8+ T cells and dissect their immunosuppressive interactions, we developed the SCLC_TR signature, which provided a robust framework for distinguishing reactive from non-reactive T cells. This signature not only outperformed established external signatures such as NeoTCR8 and TR_30 in the SCLC dataset but also demonstrated robust performance in PDAC, another external cold tumor dataset^29^. These findings suggest that SCLC_TR may be particularly suited in general to cold TMEs characteristic of PDAC and SCLC tumors. However, further studies are required to validate its performance in additional datasets derived from similar cold TMEs.

The SCLC_TR signature identified key pathways, such as the *CXCL13-CXCR5* axis linked to TLS formation and *TNFSF9-TNFRSF9* axis associated with T-cell activation, both relevant to anti-tumor immunity. These pathways have been implicated in promoting effective immune responses across various cancers, including colorectal cancer and NSCLC.^53,55^ Notably, *CXCL13*, a marker for TLS formation and B-T cell crosstalk, has emerged as a key indicator of favorable prognosis and therapeutic responsiveness and is consistently included in all other described neoantigen or tumor-reactive signatures, further highlighting its central role in tumor immunity.^25,27–29,52^ Strikingly, mature TLS formation could be detected in a patient of an independent cohort underlining the role of these structures in evoking T cell activity in line with recent literature.^54^

Furthermore, analysis of SCLC TME revealed co-dependency of tumor-reactive T cells, CD4+ T_reg_ cells and myeloid cells suggesting multiple direct and indirect immunosuppressive mechanisms. Upregulation of pathways such as *CD80/CD86-CTLA4*, *IL1* signaling, and *TNFRSF9*-mediated activation was particularly prominent in CD4+ T_reg_ cells and monocyte/macrophage populations, which are known to be able to suppress effector T-cell activity and sustain immunosuppressive networks, thereby promoting tumor progression.^60,78^ For instance, the IL1 signaling pathway amplifies T_reg_ cell activity, dampens effector T cell responses, and sustains chronic inflammation, ultimately contributing to tumor progression.^73–75^ Similarly, the *CD80/CD86-CTLA4* axis, typically observed in APCs, interacts with CTLA4 on CD4+ T_reg_ cells, enhancing their suppressive function and limiting effector T cell activation, a well-established immune checkpoint mechanism in cancer.^63^ The *TNFRSF9* (CD137) pathway exhibits a dual role in immune regulation. While CD137 signaling through pDCs or monocytes/macrophages can enhance T cell activation and survival, its persistent engagement within the TME by ligands such as *TNFSF9* may drive T cell exhaustion, further diminishing anti-tumor immunity.^61^ Additionally, the *ICOS-ICOSL* interaction supports CD4+ T_reg_ cell survival and expansion within the TME, with high *ICOS* expression on CD4+ T_reg_ cells linked to immune evasion and poor prognosis in multiple cancer types.^69^ Collectively, these mechanisms likely counterbalance anti-tumor T cell responses and exacerbate the dysfunction of tumor-reactive CD8+ T cells, further limiting their therapeutic potential.^37,65,70,71^ A deeper understanding and validation of the immunosuppressive factors could further enhance immune checkpoint inhibition which is already administered to SCLC patients, but only with marginal benefit and lead to new strategies to unleash the anti-tumor potential of tumor-reactive T cells, as targeting these pathways presents an opportunity to disrupt immune suppression and reinvigorate anti-tumor immunity. For instance, selective targeting of *IL1*-mediated signaling could effectively mitigate chronic inflammation, which supports monocyte/macrophage-driven immunosuppression and immune evasion.^73^ Furthermore, the heterogeneity in TCR reactivity and the role of immune escape mechanisms, including reduced antigen presentation, e.g. through HLA class I downregulation as seen for one of our 3 SCLC patients, and clonal diversification in metastatic sites, may pose additional challenges that highlight the need to tailor immunotherapeutic strategies.^1,2,79,80^ Further investigation is required to elucidate immune escape mechanisms and their dynamic evolution during disease progression and therapeutic resistance, which is critical for the development of more effective immunotherapeutic strategies in SCLC.

Integrating these insights into combinatorial therapy approaches engaging multiple immune components and potentially integrating multiple treatment modalities like immune checkpoint inhibitors, TCR-engineered T cells, small molecule inhibitors, cytokines, and strategies to enhance antigen presentation will ultimately lead to a more effective anti-tumor response.^30,76,81–85^ Following such potential next-generation immunotherapies, when further restraints of tumor-reactive CD8+ T cells are lifted, the prognostic potential of our SCLC_TR signature may be even more pronounced.

In summary, this study provides a comprehensive characterization of the immune landscape, centered on tumor-reactive T cells in SCLC, challenging the notion of an immune-cold tumor. By providing means to robustly identify tumor-reactive T cells and to elucidate their transcriptional states and co-stimulatory and co-inhibitory pathways, and the challenges posed by immune escape, our findings pave the way for innovative immunotherapeutic strategies. Future studies should focus on refining tumor-reactive T cell signatures and assess their applicability for different immune-cold tumor types and SCLC subtypes, optimizing TIL expansion protocols, and exploring combination therapies to overcome the barriers imposed by SCLC’s cold and immunosuppressive TME. Additionally, the development of predictive biomarkers for patient stratification and therapeutic response will be critical for the successful translation of these findings into clinical practice. These efforts hold promise for advancing the treatment of SCLC and improving outcomes for patients with this aggressive malignancy.

### Limitations of the study

While this study provides a comprehensive single-cell analysis of the SCLC immune landscape, several limitations should be acknowledged. Our TCR screening approach relies on the establishment of autologous tumor cell lines derived from PDX-models, which is time-intensive, and may not be feasible for all cases, underscoring the need for more rapid and scalable functional screening methodologies. Furthermore, as our single-cell profiling, focused on the infiltrating immune compartment in SCLC, did not capture non-immune tumor and stromal compartments, the direct assessment of their contributions to immune regulation was limited. For instance, in light of the crucial role of TLS, interaction with high endothelial venules (HEV) known to be involved in TLS formation, could be dissected further.^86^

## Acknowledgments

We thank A. Henrich and F. Durak for excellent technical assistance during sequencing; A. Agaci for faithful support in sample acquisition and processing; J. Levin for dedicated support in the cell culture; M. Graessl for committed support in cloning; R. Gudimella for supporting the survival analysis; A. Cortini for support in TCR sequence extraction and discussions; L. Bogler for support in off-target TCR screening, K. Wlotzka for supporting IncuCyte experiments. We are grateful to F. Vascotto for fruitful discussions. This work was supported by the German Research Foundation (DFG) through funding provided by a collaborative research center grant on small cell lung cancer (CRC1399, project-ID 413326622). R.K.T is also supported by the German Federal Ministry of Education and Research (BMBF, e:Med consortium InCa, grant 01ZX1901A and 01ZX2201A), by the German state of North Rhine-Westphalia EFRE initiative (EFRE-0800397) and from “Netzwerke 2021”, an initiative of the Ministry of Culture and Science of the German state of North Rhine-Westphalia for the CANTAR project.

## Declaration of generative AI-assisted technologies in the manuscript preparation process

During the preparation of this work the author used ChatGPT and Perplexity in order to enhance the text’s grammar, wording and readability. After using this tools, the authors reviewed and edited the content as needed and take full responsibility for the content of the published article.

## Resource availability

### Lead contact

Further information and requests for resources and reagents should be directed to and will be fulfilled by the lead contact, Laura Kolb

### Materials availability

We will share unique reagents generated in this study upon request. Material transfer agreements will be necessary to obtain reagents.

### Data and code availability

Raw single-cell RNA-seq, WES and RNAseq data will have been deposited at ENA and publicly available as of the date of publication. Preprocessed single-cell RNA-seq data will have been deposited at Zenodo and publicly available as of the date of publication. Scripts used to process the data and generate the figures will be available upon request as of the date of publication.

### Experimental model and study participant details

#### Human lung tumor specimen

This study was approved by the Institutional Review Board (IRB) of the University of Cologne and conducted in accordance with the Declaration of Helsinki. Written informed consent was obtained from all patients prior to tissue collection.

### Method details

#### Sample collection

A total of 22 tumor samples were collected from 20 patients with histologically confirmed lung cancer, either through diagnostic fine-needle biopsies or therapeutic surgical resections, under IRB-approved protocols coordinated through a network of academic hospitals and clinical research institutions. All samples underwent pathological evaluation by at least 2 board-certified pulmonary pathologists. Histomorphology classification was performed using haematoxylin and eosin (H&E) staining, and immunohistochemical (IHC) staining was used to aid in the differentiation of neuroendocrine and non-neuroendocrine subtypes. Tumors were classified according to World Health Organization (WHO) criteria as SCLC, NSCLC, or LCNEC. Although all specimens were primary tumors, lymph node metastases were also included in a subset of cases (Patients 1, 3, and 6) to provide additional insights into metastatic progression and tumor microenvironment heterogeneity.

#### Patient-derived xenograft cell line generation

Fresh tumor samples were acquired as fine-needle biopsies. For cell line generation, the tumor tissues were engrafted onto immune-compromised mice (NSG mice) to establish PDX, allowing functional validation of tumor-reactive TCRs in co-culture assays and genomic studies. All animals were housed in a specific pathogen-free facility under ambient temperature and humidity while maintaining a 12/12 h light/dark cycle. Animal experiments were approved by, and conducted in accordance with, the regulations of the local animal welfare authorities (State Agency for Nature, Environment, and Consumer Protection of the State of North Rhine-Westphalia).

#### Culture of tumor cells derived from patient-derived xenografts (PDXs)

PDX tumor cell lines were established from SCLC tissue samples obtained from patient 5, 6, and 7. Tumor fragments were engrafted subcutaneously into immunodeficient mice (e.g., NSG or NOD SCID strains), and xenografts were harvested once tumors reached an appropriate size. Fresh tumor tissues were dissociated into single-cell suspensions, and PDX-derived cell lines were subsequently cultured in a defined serum-free medium optimized for the propagation of epithelial tumor cells. Cells were maintained in Advanced DMEM/F12 (Gibco, Cat# 12634028) supplemented with 1× B-27 supplement (Gibco, Cat# 17504044), 10 mM HEPES (Gibco, Cat# 15630080), 1× GlutaMAX (Gibco, Cat# 35050061), and 100 U/mL Penicillin and 100 µg/mL Streptomycin (Gibco, Cat# 15140122) to support cell viability and minimize microbial contamination. To support growth and self-renewal, cultures were further enriched with 50 ng/mL R-Spondin 1 (PeproTech, Cat# 120-38-250UG9), 100 ng/mL FGF10 (PeproTech, Cat# AF-100-26-25UG), and 25 ng/mL FGF7 (PeproTech, Cat# 100-19-10UG), key mitogens known to promote lung epithelial cell expansion. Additionally, small molecule inhibitors were included to enhance survival and inhibit unwanted differentiation: 500 nM A83-01 (Sigma-Aldrich, Cat# SML0788), a TGF-β receptor inhibitor, and 5 µM Y-27632 (AbMole, Cat# M1817), a ROCK inhibitor that prevents apoptosis during cell stress and dissociation. To prevent contamination, all cultures were supplemented with 50 µg/mL Primocin (InvivoGen, Cat# ant-pm-05), an antibiotic formulation active against bacteria, mycoplasma, and fungi. Cells were cultured in a humidified incubator at 37 °C with 5% CO₂ and were passaged approximately once per week, depending on confluence and proliferation rate. Medium changes were performed every 2–3 days to maintain optimal nutrient levels and growth factor activity. Cells were monitored regularly for morphology, viability, and contamination, and only early-passage PDX cultures were used for functional assays to preserve phenotypic fidelity.

#### Cryopreservation of tumor tissue using slow freeze protocol

Tumor tissues were cryopreserved using a controlled slow-freeze protocol to preserve viability and structural integrity for future analyses. Upon arrival, tissue specimens in transport medium were transferred into sterile 15 cm Petri dishes (Corning, Cat# 430167) using autoclaved forceps. The tissue was gently placed onto a precision balance (e.g., Sartorius Entris series), and its weight was recorded. A photograph was taken of the tissue with the scale underneath for documentation and sample traceability. For decontamination, 3 adjacent wells of a 6-well plate (Corning, Cat# 3516) were filled with 5 mL of Washing Medium, which consisted of XVIVO medium (Lonza, Cat# 04-418Q) supplemented with Penicillin-Streptomycin (Gibco, Cat# 15140122; 10,000 IU/mL; added at 500 µL per 50 mL) and Amphotericin B (Gibco, Cat# 15290026; 0.5 mL of a 250 µg/mL stock per 50 mL). The tissue was sequentially washed by transferring it from one well to the next, gently swirling with a sterile pair of autoclaved forceps at each step to remove residual media and contaminants. After washing, the tissue was transferred to a new sterile 15 cm Petri dish containing 2–3 mL of Dissection Medium, composed of XVIVO supplemented with 2% human serum albumin (HSA; Grifols, Cat# 081618701), Penicillin-Streptomycin, and Amphotericin B as previously described. Using sterile surgical scalpels (Feather, No. 11, Cat# 05-080-003, Fisher Scientific), the tissue was finely chopped into approximately 5 mm² fragments. The number of fragments was recorded, and a second image was taken to log the condition and quantity of the material. One tissue fragment was then transferred into each cryopreservation tube (Nalgene, Cat# 5000-0020), and its source location within the original tissue was documented for reference. To each tube, 1 mL of freshly prepared Freezing Medium—composed of fetal calf serum (FCS; Gibco, Cat# 10082147) supplemented with 10% dimethyl sulfoxide (DMSO; Sigma-Aldrich, Cat# D2650)—was added. The cryotubes were tightly sealed and immediately placed in a Mr. Frosty freezing container (Thermo Fisher Scientific, Cat# 5100-0001) pre-filled with isopropanol and stored at −80 °C to allow gradual cooling at approximately −1 °C per minute. After incubation for a minimum of 16 hours and no more than 96 hours at −80 °C, cryovials were transferred to vapor-phase liquid nitrogen storage tanks (approximately −196 °C) for long-term preservation.

#### DNA & RNA extraction from cells & fresh frozen tissue

According to the manufacturers protocol, DNA from cells and fresh frozen tissue was extracted using Qiagens DNeasy Blood & Tissue Kit. For RNA extraction Qiagens RNeasy Mini Kit for total RNA was used. Quantity and quality were assessed using the Qubit 3 fluorometer – RNA HS Assay kit (Invitrogen) for RNA or the 1X dsDNA HS Assay Kit (Invitrogen) for DNA and the Nanodrop2000. RNA quality (RIN) was additionally determined using the Agilent Bioanalyzer 2100 system using the RNA Pico kit.

#### NGS library preparation

Whole exome libraries were prepared in duplicates with an input of 100 ng total RNA or genomic DNA each. All samples were fragmented, end repaired and adenylated – the RNA samples were primed and reverse-transcribed after fragmentation first. An appropriate single index eight-nucleotide NEXTFLEX DNA barcode (Perkin Elmer) was ligated for the pre-amplification of the library which was done using the KAPA Hyper Prep kit (Roche). Subsequently, target regions were hybridized to biotinylated baits (SureSelectXT Human All Exon v6 and SureSelect XT Reagent kit, Agilent) and isolated using streptavidin-coated magnetic beads (Invitrogen). The post-capture library was then amplified with the KAPA Library Amplification kit (Roche). All intermediate and final library purification steps were carried out using AMPure XP beads (Beckman Coulter).

All QC steps for library preparation were performed using the Qubit 3 fluorometer (1X dsDNA HS Assay Kit, Invitrogen) for quantitative analysis and the Agilent Bioanalyzer 2100 system (High Sensitivity DNA Kit, Agilent) for qualitative analysis.

#### Exome and RNA Sequencing

All libraries were sequenced in paired-end mode (2 x 50 nt) on an Illumina NovaSeq 6000 instrument resulting in around 75 million distinct sequencing reads per RNA Exome Capture Library or 150 million reads per DNA Exome Capture Library.

#### Mutation calling

Somatic mutations were detected as described previously^87,88^ with an in-house pipeline. In short, DNA reads were aligned to the hg19 reference genome using BWA (v0.7.11)^89^ and duplicated reads were removed with Picard (v1.110) (https://broadinstitute.github.io/picard/). High-confidence single nucleotide variants were detected by an in-house propriety software.

#### RNA expression analysis

For RNA expression analysis, RNA-seq reads were aligned to the hg19 reference genome using STAR (v2.4.2a).^90^ Transcripts were quantified in FPKM with sailfish (v 0.7.6).^91^ Gene expression was approximated by the sum of transcript expression of all transcripts related to a gene.

#### Sample processing for single-cell transcriptomic and V(D)J sequencing

Following initial tumor dissection, samples were first mechanically processed using sterile scalpels (Feather, No. 11, Cat# 05-080-003, Fisher Scientific) and then finely minced into 1–2 mm fragments on a sterile Petri dish (Corning, Cat# 430167) under aseptic conditions. The fine minced fragments were transferred into a 5 mL tube and incubated in 1 mL of pre-warmed digestion buffer (DB), consisting of Advanced DMEM/F12 (Gibco, Cat# 12634010), 75 µg/mL Liberase TL (Roche, Cat# 5401020001), and 2 µg/mL DNase I (Roche, Cat# 11284932001). Gentle mixing was performed using a wide bore pipette tip (Axygen, Cat# TF-1000-WB-R-S), followed by incubation at 37 °C. During digestion, samples were gently shaken at 5 and 10 minutes and mixed by using a wide-bore pipette tip at 15 and 25 minutes. After each mixing step, the tissue was allowed to settle for 1–2 minutes. Subsequently, 300 µL of partially digested material was collected and transferred to a pre-chilled 50 mL tube containing 10 mL of collection buffer (CB) and topped with a 70 µm cell strainer (Corning, Cat# 352350). The strainer was rinsed with an additional 300 µL of CB (Advanced DMEM, Gibco, Cat# 12491023, supplemented with 1% heat-inactivated FBS, Gibco, Cat# 10082147) to recover cells. This collection step was repeated every 10 minutes using progressively narrower cut tips, transitioning to an uncut tip as the tissue softened. From the third round onward, 300 µL of fresh digestion buffer was added after each collection step to maintain enzymatic activity. The digestion process continued until only residual extracellular matrix (ECM) remained, typically after approximately 60 minutes. The final digestion mixture was passed through the 70 µm strainer using the plunger of a sterile 10 mL syringe (BD, Cat# 302995). The original digestion tube was rinsed with 10 mL of cold CB, which was also passed through the same strainer. The resulting single-cell suspension was centrifuged at 300 ×g for 8 minutes at 4 °C (Eppendorf 5810R centrifuge). The cell pellet was resuspended in 200 µL of flow cytometry buffer (1x PBS (Ca2+/Mg2+ free), 0.5 mM EDTA, 2% FCS) for downstream staining and sorting or frozen in medium consisting of FCS (Gibco, Cat# 10082147) supplemented with 10% DMSO (Sigma-Aldrich, Cat# D2650).

#### Preparation and sorting of CD45⁺ single-cell suspensions

Single-cell suspensions were prepared for fluorescence-activated cell sorting (FACS) to isolate CD45⁺ immune cells using a two-step staining protocol. Cells obtained from the prior tissue dissociation step or post thawing the frozen single-cell suspensions were transferred into low-binding 1.5 mL microcentrifuge tubes (Eppendorf DNA LoBind, Cat# 0030108051) and centrifuged at 300 ×g for 8 minutes at 4 °C. The cell pellet was resuspended in 1 mL of FACS buffer, composed of phosphate-buffered saline (PBS; Gibco, Cat# 10010023) supplemented with 2% fetal bovine serum (FBS; Gibco, Cat# 10082147) and 0.5 mM EDTA (Thermo Fisher Scientific, Cat# 15575020). For viability staining, the supernatant was discarded, and cells were incubated with Fixable Viability Dye eFluor 780 (Thermo Fisher Scientific, Cat# 65-0865-14) diluted 1:1167 (0.1 µL in 149.9 µL FACS buffer per sample) in the dark at 4 °C for 20 minutes. Following incubation, cells were washed by centrifugation at 300 ×g for 8 minutes, the supernatant was discarded, and the pellet was resuspended in 200 µL of FACS buffer. Surface staining was then performed using a panel of fluorochrome-conjugated monoclonal antibodies, each diluted in FACS buffer to a final volume of 100 µL per tube. The staining panel included: CD45 PE-Cy7 (BD Biosciences, Cat# 557748; 1 µL, 1:100), CD4 PE (BioLegend, Cat# 317408; 2 µL, 1:50), CD3 FITC (BioLegend, Cat# 300406; 2 µL, 1:50), CD8 APC (BD Biosciences, Cat# 561421; 1 µL, 1:100), CD16 PerCP-Cy5.5 (BioLegend, Cat# 302028; 2 µL, 1:50), CD56 BV786 (BD Biosciences, Cat# 564058; 2 µL, 1:50), CD137 BV421 (BioLegend, Cat# 309826; 1 µL, 1:100), and CD274 (PD-L1) BV510 (BD Biosciences, Cat# 564715; 2 µL, 1:50). An unstained control tube received 100 µL of FACS buffer alone. Samples were incubated in the dark at 4 °C for 20 minutes, followed by washing via centrifugation at 300 ×g for 8 minutes. The pellet was resuspended in 200 µL of FACS buffer with gentle pipetting. Final stained samples were transferred to pre-labeled 5 mL polystyrene FACS tubes (Falcon, Cat# 352008), matched to their respective antibody panels, and kept on ice or at 4 °C prior to flow cytometric analysis or cell sorting using a BD FACSAria or equivalent instrument.

#### Preparation and sequencing of single-cell transcriptomic and V(D)J libraries

Live CD45+ cells were sorted into 1X PBS with 0.04 % BSA (PBS-BSA) into low-binding 1.5 mL microcentrifuge tubes (Eppendorf DNA LoBind, Cat# 0030108051). Final cell suspensions were processed according to 10X Genomics Chromium Single Cell 5’ Reagent Guidelines (Chromium Single Cell V(D)J Reagent Kits v1.1 or Chromium Single Cell V(D)J Amplification Kits (v2.0)). Quantification of cDNA and final libraries were performed using Qubit dsDNA HS Assay Kit (Life Technologies) and High-sensitivity DNA Chips (Agilent). Final libraries were diluted to 200 pM, pooled and loaded on an Illumina NovaSeq 6000 SP, S1or S2 flow cell depending on experiment using the NovaSeqTM 6000 SP/S1/S2 SBS Cartridge (100 cycles) and NovaSeqTM 6000 SP/S1/S2 Buffer Cartridge. Libraries were sequenced using a NovaSeqTM 6000 with the following read lengths: 26 cycles (Read 1), 8 cycles (i7 Index) and 90 cycles (Read 2) for single index for Patients 1, 6 and 7. For all other patients where dual index settings were used (28 cycles (Read 1), 10 cycles (i7 Index), 10 cycles (i5Index) and 90 cycles (Read 2)).

#### Single-cell RNA sequencing raw data processing and analysis

The 10x Genomics Cell Ranger pipeline (versions 6.0.2 or 7.0.0 (with parameter --include-introns=false), depending on sequencing batch) was used for processing scRNA-seq and V(D)J data. The cellranger mkfastq function was employed to demultiplex raw base call (BCL) files into FASTQ files. Reads were aligned to the GRCh38-3.0.0 human reference genome (refdata-cellranger-GRCh38-3.0.0) using the cellranger count function. Across samples, captured cell numbers ranged from approximately 205 to 11,812 per library, with ∼83,000 to 189,000 reads per cell and a median of ∼1200 to 3,625 genes per cell (Supplementary Table S1). For Patient 3, the cellranger aggr pipeline was run on the duplicates to create a single feature barcode matrix per site. For V(D)J analysis, paired TCR sequences were reconstructed using the cellranger vdj function. FASTQ files were aligned to the appropriate reference (refdata-cellranger-vdj-GRCh38-alts-ensembl-5.0.0) to annotate full-length, paired V(D)J sequences. Contigs were filtered to retain only high-confidence, productive chains. Each T cell was assigned a clonotype based on identical CDR3 nucleotide sequences and V(D)J gene usage. Clonotype tables were used for downstream repertoire analysis, including clonal expansion assessment, diversity analysis, and integration with transcriptomic profiles via barcode-matching. Captured cell numbers with TCRs ranged from approximately 1702 to 4718 per sample library, with 35,138 to 112,000 reads per cell and a median of ∼1200 to 1551 genes per cell for the scRNA-seq data. For downstream scRNA-seq analysis, Seurat v4.2.0 (with R v4.1.0) was used following the standard Seurat integration workflow (https://satijalab.org/seurat/). Cells with fewer than 200 detected genes, or greater than 3 median absolute deviations (MADs) above the median for mitochondrial gene content or gene counts, were excluded. Expression matrices were log-normalized and transformed, with 2,000 variable features selected for downstream analysis. Each sample was processed individually until integration, which was performed using canonical correlation analysis (CCA) and anchor-based alignment. A linear transformation (scaling) was applied to all cells, and principal component analysis (PCA) was performed. Dimensionality reduction and visualization were conducted using Uniform Manifold Approximation and Projection (UMAP), with clustering based on the top 17 principal components at a resolution of 1.

Cluster annotation was performed using canonical markers and differentially expressed genes, with low-quality clusters—characterized by low RNA content, high mitochondrial reads, ambient RNA contamination, or doublet signals—excluded from further analysis. Additional doublets were removed using scDblFinder (version 1.13.5). To improve granularity, several rounds of subclustering were conducted on defined populations including myeloid cells and combined B and T lymphocytes. Final refined annotations were transferred back to the integrated Seurat object for comprehensive analysis and visualization. Cell annotations were based on canonical markers previously reported in literature. From the CD4+ T cell clusters, ‘CD4+ TH1’ showed expression of Type I T helper cells markers including CD4, IFNG and TBX21.1, 2 ‘CD4+ Treg cells’ cluster was marked by the expression of FOXP3, IKZF2, IL2RA, CTLA4, TIGIT.3 An additional ‘CD4+ effector’ cluster was identified which lacked the expression of FOXP3 and IFNG but expressed other effector genes including IL22, RORC, IL2RA. The clusters ‘Naïve/memory T cells’, ‘CD4+ naïve/memory T cells’ and ‘CD8+ effector memory T cells’ were characterised by high expression of SELL, TCF7, CCR7, LEF1. ‘CD8+ effector memory T cells additionally express GZMK, GNLY. CD8+ T cells effector states were marked by expression of several cytotoxicity, exhaustion and proliferation related genes including GZMB, GNLY, PRF1, GZMK, GZMH, LAG3, TNFRSF9, TIGIT, TOX, TOP2A and BIRC5. NK cells cluster express NCAM1, NCR1, NCR3, KLRC1 which have been reported as activating and inhibitory receptors in tumour-infiltrating NK cells. The B cells cluster is characterised by the expression of CD79A, MS4A1, JCHAIN, IGHA1 genes. The myeloid cells co-expressed CD68, CSF1R, CD86. Additionally, non-classical monocytes cluster expressed FCGR3A, CDKN1C, LILBR2 and CXCL10. The Monocytes/macrophage cluster expressed CD14, MRC1, APOE, MACRO, and CXCL9. The DC2 cluster showed expression of FCER1A, CD74, XCR1, CLEC9A, CD1C and CLEC10A and the pDCs were marked by the presence of LILR4A, SERPINF1 and IL3RA.

#### Single-cell T Cell Receptor (scTCR) repertoire analysis

TCR V(D)J repertoire analysis was performed using the scRepertoire (v2.0.7) and Immunarch (v0.6.8) R packages, focusing exclusively on αβ TCR chains (TRA and TRB). Productive, high-confidence paired TCR sequences were derived from cellranger vdj outputs and matched with single-cell transcriptomic data using shared barcodes. Clonotypes were defined by exact CDR3 nucleotide sequences in combination with V and J gene usage. Cells with paired TRA and TRB chains were retained for analysis, and clonotype annotations were integrated into the corresponding Seurat object using the combineExpression() function. To explore clonal expansion patterns, clones were categorized into expansion bins (e.g., singletons, small [2–5 cells], moderate [6–20 cells], large [>20 cells]) and visualized across UMAP embeddings. The clonalOccupy() function was used to assess the relative contribution of dominant clones across transcriptionally defined T cell clusters, providing insight into clonal skewing and dominance within phenotypically distinct subsets. Alluvial plots (alluvialClonotypes()) were generated to track clonal transitions across sample groups, timepoints, or T cell states, illustrating the lineage relationships and stability of clonotypes over experimental conditions. Clonal abundance was profiled using frequency bar plots, and clonal homeostasis distributions were quantified to evaluate the proportion of expanded versus unexpanded clones across samples. Clonal diversity was assessed using the Shannon entropy and inverse Simpson indices and overlap between samples or groups was computed using both scRepertoire and Immunarch, with metrics such as Jaccard and Morisita indices. V and J gene usage heatmaps were also constructed using Immunarch to compare gene segment preferences across samples or clusters.

#### Prioritization of candidate tumor-reactive TCRs

To prioritize tumor-reactive TCR clonotypes, we employed a two-step signature scoring approach based on the NeoTCR8 gene signature. First, gene set variation analysis (GSVA) was used to evaluate the enrichment of the NeoTCR8 signature across transcriptionally defined CD8⁺ T cell subsets. GSVA was performed using the gsva() function from the GSVA R package (v1.46.0) on log-normalized expression data extracted from the integrated Seurat object. This analysis enabled the identification of CD8⁺ clusters with the highest overall enrichment of tumor-reactive transcriptional features. GSVA scores were displayed in a heatmap, and high-scoring clusters were chosen for further investigation.Next, within the most enriched subset—CD8⁺ effectors_2—we applied UCell scoring to evaluate NeoTCR8 signature expression at the resolution of individual cells. UCell scores were implemented via the UCell R package (v2.2.0). The NeoTCR8 gene set was used as input to the ScoreSignatures_UCell() function, and resulting scores were assigned to each CD8⁺ effectors_2 cell. TCR clonotypes within this population were then ranked based on the average UCell score of cells expressing each clonotype, allowing prioritization of putatively tumor-reactive TCRs for functional validation.

#### DNA template manufacturing for TCR expression

Patient-specific variable regions of TCRα and TCRβ chains were synthesized and cloned into custom expression vectors encoding the corresponding human TCR constant regions (Twist Bioscience, Cat# PN: custom synthesis). Each variable region was inserted into its respective plasmid backbone to enable efficient *in vitro* transcription and expression of the full-length TCR. Amplification of TCRα and TCRβ inserts was performed via polymerase chain reaction (PCR) using high-fidelity Q5® Hot Start High-Fidelity DNA Polymerase (New England Biolabs, Cat# M0493S). Reactions were carried out with the following primers:

- Forward primer: 5′-GTTCATCTGCACCACCGGCAAGCTG-3′
- Reverse primer: 5′-(T)₁₂₀AGCGACGCGG-3′

PCR reactions were run under standard cycling conditions optimized for specificity and high-yield amplification. Post-PCR products were purified using the Monarch® PCR & DNA Cleanup Kit (Spin Columns) (New England Biolabs, Cat# T1030L), following the manufacturer’s instructions. Purified DNA was eluted in nuclease-free water (Thermo Fisher Scientific, Cat# AM9937) and quantified using a NanoDrop spectrophotometer (Thermo Fisher Scientific, Cat# ND-ONE-W). For scalable, high-throughput template generation, all liquid handling steps including PCR setup, reaction clean-up, and normalization were automated using the Biomek i7 Automated Workstation (Beckman Coulter, Cat# B24402). The platform was programmed according to manufacturer protocols and operated in conjunction with PCR-compatible 96-well plates (Bio-Rad, Cat# HSP9601) and magnetic bead–based purification systems, when applicable. This workflow provided high-quality, linearized DNA templates suitable for subsequent *in vitro* transcription and T cell electroporation applications.

#### Generation of *in vitro* transcribed (IVT) RNA

PCR-amplified linear DNA templates encoding full-length TCRα and TCRβ sequences were used for *in vitro* transcription (IVT) of RNA. Transcription reactions were performed using the HiScribe™ T7 Quick High Yield RNA Synthesis Kit (New England Biolabs, Cat# E2050S) in accordance with the manufacturer’s protocol. Each reaction was assembled using 1–2 µg of purified DNA template and incubated at 37 °C for 2–4 hours to maximize RNA yield. Following IVT, RNA was treated with DNase I (included in the kit) to eliminate residual template DNA. Transcribed RNA was purified using Monarch® RNA Cleanup Kit (50 µg capacity, Spin Columns) (New England Biolabs, Cat# T2040S) to remove enzymes, unincorporated nucleotides, and other reaction components. Eluted RNA was resuspended in nuclease-free water (Thermo Fisher Scientific, Cat# AM9937). RNA concentration was quantified using the NanoDrop™ 2000 Spectrophotometer (Thermo Fisher Scientific, Cat# ND-2000), and RNA integrity was assessed using the Fragment Analyzer System (Agilent, formerly Advanced Analytical, Cat# DNF-472-0500 for RNA Analysis Kit). Samples were loaded according to the manufacturer’s protocol to verify full-length transcript production and to ensure the absence of degradation products. Only RNA samples meeting stringent quality criteria—including correct size and high integrity (RQN>8)—were used for downstream electroporation into CRISPR-edited T cells.

#### Isolation, CRISPR editing, and electroporation of primary human CD8⁺ T Cells

Primary human CD8⁺ T cells were isolated from fresh buffy coats obtained from healthy donors through the blood donation service at the University Medical Center Mainz, in accordance with institutional ethical guidelines. Peripheral blood mononuclear cells (PBMCs) were enriched using the StraightFrom® Whole Blood CD8 MicroBead Kit (Miltenyi Biotec, Cat# 130-090-878), following the manufacturer’s protocol for positive magnetic selection. Isolated CD8⁺ T cells were cultured in X-VIVO 15 medium (Lonza, Cat# 02-053Q) supplemented with 50 U/mL recombinant human IL-2 (PeproTech, Cat# 200-02-1MG).

To enable knockout of endogenous T cell receptor (TCR) expression, CD8⁺ T cells were first pre-activated for 3 days using TransACT™ CD3/CD28 nanomatrix (Miltenyi Biotec, Cat# 130-111-160) at the manufacturer-recommended concentration, in X-VIVO 15 medium supplemented with 100 U/mL IL-2. Following activation, cells were rested for 24 hours in IL-2–containing medium prior to electroporation. For CRISPR-mediated knockout of the endogenous TCR, up to 15 × 10⁶ pre-activated T cells were resuspended in 125 µL of fresh X-VIVO 15 medium and mixed with a ribonucleoprotein (RNP) complex containing 2.5 µg of chemically synthesized single-guide RNAs (sgRNAs) targeting the TRAC locus (sequence: AGAGUCUCUCAGCUGGUACA) and the TRBC locus (sequence: GGAGAAUGACGAGUGGACCC), along with 5 µg of *in vitro*–transcribed Cas9 mRNA (TriLink BioTechnologies, Cat# L-7206 or equivalent). Electroporation was performed using the ECM 830 electroporation system (BTX Harvard Apparatus, Cat# 45-2052) set to 225 V for 5 milliseconds in a 2 mm cuvette (BTX, Cat# 45-0124). Immediately after electroporation, cells were transferred into pre-warmed X-VIVO 15 medium supplemented with 100 U/mL IL-2 and incubated at 37 °C in a humidified 5% CO₂ incubator. After confirming efficient TCR knockout by flow cytometry using anti-human TCRα/β (BioLegend Cat# 306718), anti-human CD8 (BioLegend Cat# 344745), and Fixable Viability Dye eFluor 780 (Thermo Fisher Cat# 65-0865-14), cells were rested for an additional 24 hours. Subsequently, CRISPR-edited T cells were resuspended at a concentration of 2 × 10⁶ cells in 125 µL X-VIVO 15 medium and electroporated with 10 µg of *in vitro*–transcribed RNA encoding the respective TCRα and TCRβ chains. The same electroporation parameters were used (225 V, 5 ms pulse, ECM 830 system). Transfected cells were returned to culture in IL-2–supplemented X-VIVO 15 and incubated for 16–18 hours prior to flow cytometric analysis and functional assays.

#### Disintegration of tumor spheroids

Tumor spheroids used as target cells in cytotoxicity or ELISpot assays were disaggregated into single-cell suspensions using a two-round enzymatic digestion protocol. Spheroids were first harvested using a 10 mL serological pipette and transferred into a 15 mL conical tube (Falcon, Cat# 352096). The suspension was centrifuged at 300 ×g for 15 seconds, stopping the centrifuge immediately upon reaching the set speed to preserve spheroid integrity. The supernatant, typically cloudy with residual medium, was carefully aspirated, ensuring the spheroids remained at the bottom. To initiate dissociation, 2 mL of pre-warmed Accutase® Cell Detachment Solution (Innovative Cell Technologies, Cat# AT104) was added to the pellet, followed by incubation at 37 °C for 20 minutes. At the 10-minute mark, the suspension was gently mixed by pipetting to facilitate enzymatic access. The partially dissociated mixture was then passed through a 70 µm MACS SmartStrainer (Miltenyi Biotec, Cat# 130-098-462) into a fresh 15 mL Falcon tube to collect single cells from the first digestion round (Round 1). Cells were centrifuged at 300 ×g for 8 minutes at room temperature, the supernatant was aspirated, and the pellet was resuspended in ice-cold phosphate-buffered saline (PBS) (Gibco, Cat# 10010023).

To maximize cell yield, the strainer containing residual clumps was inverted into a new Falcon tube, and the retained tissue was subjected to a second digestion round. An additional 2 mL of Accutase was added directly to the flipped strainer, and incubation continued at 37 °C for another 5–10 minutes. Cells from Round 2 were then centrifuged and resuspended in PBS as before. Cell numbers from each round were manually quantified using the C-Chip disposable counting chamber (NanoEnTek, Cat# DHC-N01), and viability was assessed using Erythrosine B (Logos Biosystems, Cat# L13002). If sufficient cell numbers were obtained from first round, the second fraction was not combined. Final target cell suspensions were adjusted to a concentration of 5 × 10⁵ cells/mL (equivalent to 50,000 cells per 100 µL) in T cell culture medium to avoid pre-activation or proliferation of co-cultured immune cells.

#### Interferon-γ enzyme-linked immunospot (ELISpot) assay

To evaluate antigen-specific T cell responses, interferon-γ (IFN-γ) ELISpot assays were performed using co-cultures of tumor cells and TCR-transfected CD8⁺ T cells. Briefly, 5 × 10⁴ patient-derived tumor cells were seeded per well in ELISpot plates pre-coated with anti-IFN-γ capture antibody (Millipore, Cat# MSIPS4510), which had been precoated with anti-human IFN-γ monoclonal antibody (clone 1-D1K) (Mabtech, Cat# 3420-3-250). After tumor cell adherence, 1 × 10⁵ CD8⁺ T cells—previously electroporated with *in vitro*–transcribed TCR mRNA and rested for 16–18 hours—were added directly to each well for co-culture. Plates were incubated for 20 hours at 37 °C in a humidified atmosphere with 5% CO₂ to allow for IFN-γ secretion in response to antigen recognition. Following incubation, cells were gently washed away with PBS, and secreted IFN-γ was detected using a biotin-conjugated anti-human IFN-γ detection antibody (clone 7-B6-1) (Mabtech, Cat# 3420-9A-1000), followed by development with BCIP/NBT-plus substrate solution (Mabtech, Cat# 3650-10), which results in visible dark blue spots at the site of cytokine release. All experimental conditions were plated in technical duplicates to ensure data reproducibility. Plates were air-dried overnight and scanned using an ImmunoSpot® Analyzer (Cellular Technology Limited, CTL) for automated spot detection and quantification. T cell reactivity was defined based on a minimum two-fold increase in the number of spots in co-culture wells compared to T cells cultured alone (background control), indicating specific IFN-γ secretion upon recognition of tumor antigens.

#### *In vitro* off-target TCR assessment

Transgenic Jurkat cells knocked out for the endogenous TCR expression and transduced to express human CD8α were generated in-house and maintained in Iscove’s Modified Dulbecco’s Medium (IMDM, Hi-Media) supplemented with 10 % fetal calf serum (FCS, Bio&SELL) and 100 µg/ml Zeocin (InvivoGen). HLA-negative K-562 cells were cultured in IMDM supplemented with 10 % FCS.

For the off-target (potential self-reactivity) assessment, 10 µg of the corresponding TCR alpha and beta chain mRNA were electroporated into 2 x 10^6^ Jurkat cells using a single pulse of 275 V and 10 ms duration using the ECM® 830 electroporation device (BTX). In parallel, 10 µg of the respective mRNA encoding patient HLA haplotypes were electroporated into 2 x 10^6^ K-562 cells using 3 pulses with 260 V and a duration of 8 ms. Cells were rested overnight and 1 x 10^5^ of both cell types were co-cultured for 6 hours in dark 96-well plates (Thermo Scientific). After incubation, the same volume of QUANTI-Luc™ 4 Lucia/Gaussia (InvivoGen) solution was added per well and secreted luciferase was measured using the Infinite M Plex microplate reader (Tecan).

#### Live imaging-based cytotoxicity assay

Real-time cytotoxicity of TCR-transfected CD8⁺ T cells was assessed using a live-cell imaging approach. Prior to co-culture, patient-derived tumor cells were labeled with 0.33 µM Incucyte® Cytolight Rapid Red Dye (Sartorius, Cat# 4706), while effector CD8⁺ T cells were labeled with 0.11 µM Incucyte® Cytolight Rapid Green Dye (Sartorius, Cat# 4705). Both cell types were incubated with their respective dyes for 20 minutes at 37 °C in serum-free medium, followed by washing with complete culture medium to remove excess dye. After labeling, 1 × 10⁴ tumor cells were seeded into ultra-low attachment 96-well plates (Costar®, Cat# CLS7007) to promote uniform spheroid formation. Tumor cells were allowed to settle for 30 minutes before the addition of TCR-transfected CD8⁺ T cells at various effector-to-target (E:T) ratios. Co-cultures were imaged using the Incucyte® SX5 Live-Cell Analysis System (Sartorius) equipped with a 4× objective lens. Time-lapse imaging was performed under standard culture conditions (37 °C, 5% CO₂), with images captured every 2 hours over the course of the assay. Fluorescent channels were used to independently track tumor (red) and T cell (green) populations, enabling real-time assessment of immune cell engagement and tumor killing. Quantitative analysis of tumor cell viability and T cell dynamics was performed using integrated Incucyte® software based on red fluorescence intensity and morphological disruption of tumor spheroids. Statistical significance between paired conditions was assessed using a two-tailed paired t-test. Unless otherwise indicated, data are presented as mean ± s.e.m.

#### Differential gene expression analysis (DGEA)

To identify genes distinguishing tumor-reactive (TR) from non-tumor-reactive (NTR) CD8⁺ T cells, differential gene expression analysis was performed using the FindMarkers() function in the Seurat R package (v4.2.0). Only CD8⁺ T cells were included in the analysis, and cells from the CD8⁺ effectors_2 cluster were excluded to avoid inclusion of clonotypes with unverified reactivity status. The analysis was conducted using the Wilcoxon rank-sum test, and genes were filtered based on a log₂ fold change threshold of >0.5 and an adjusted p-value <0.05. Shared genes expressed in both TR and NTR populations were excluded to ensure selection of tumor-reactive-specific gene features. Genes that passed these criteria were retained for further analysis as part of a putative tumor-reactive-specific transcriptional signature. Following DGEA, a tumor-reactive gene signature was derived by selecting genes upregulated specifically in the tumor-reactive CD8⁺ population. Filtering was based on expression specificity, statistical significance, and fold-change criteria as outlined above. The resulting gene set was curated and annotated for known functional associations with T cell activation, effector function, exhaustion, and immune regulation. This gene signature was used for downstream enrichment analysis and as a reference for scoring tumor-reactive states in single-cell data.

#### Ligand-receptor interaction analysis

Intercellular communication between TR/NTR CD8⁺ T cells and other immune cell types was assessed using the CellChat R package (v1.6.1). Annotated cell populations, as shown in the corresponding figure, were assigned as source and target populations for subsequent analyses. The normalized Seurat object was converted and ligand–receptor communication probabilities were computed using computeCommunProb(). Network visualizations and pathway-specific interaction maps were generated using netVisual_circle() and netAnalysis_signalingRole_network(). Predicted signaling interactions were filtered and grouped by pathway and ligand–receptor pair annotations from the CellChat database. Crosstalk among immune subsets was characterized to identify signaling axes potentially contributing to T cell activation, suppression, or differentiation.

#### Phenocycler Fusion assay

The PhenoCycler-Fusion system (Akoya Biosciences) was used to analyze tumor and immune cell populations within SCLC tumor microenvironment. Sample preparation and tissue staining were performed according to the PhenoCycler-Fusion user manual (Akoya, PhenoImager Fusion SW Version: 2.1.0). Briefly, 3 μm FFPE tissue sections were pre-processed by deparaffinization, dewaxing/rehydration, and heat-mediated antigen retrieval (Akoya, 7000017). Then, tissues were simultaneously stained with the entire barcoded-antibody panel (see Table 1). After the staining, a flow cell (Akoya, 240205) was placed on top of the slide.

For the reporter plate, unique and spectrally distinct reporters, complementary to the barcodes used in the antibody panel, were organized into groups of 3 and combined with a nuclear stain (Akoya, 7000003). Each group constituted a separate PhenoCycler cycle. The experimental protocol and reporter plate design were carried out using the PhenoCycler Experiment Designer software. The PhenoCycler run was fully automated and executed by the Controller software. PhenoCycler reporters were delivered to the tissue by the PhenoCycler instrument and detected using the Fusion slide scanner. The repetition of these cycles with different reporters allowed the visualization of the complete antibody panel on the same tissue area.

Image analysis was performed using QuPath (Version 0.5.0). Whole tissue annotation and segmentation were based on DAPI signal, with tissue segmentation conducted using the StarDist plugin. For each marker, object classifiers were independently developed using a representative training image compiled from all samples to avoid bias. Classifiers were based on characteristic features such as signal intensity patterns, subcellular localization, and morphological criteria. After segmentation, post hoc filtering excluded objects smaller than 10 µm, larger than 150 µm, and those intersecting annotation boundaries. The trained classifiers were then applied sequentially to each segmented sample. Single-cell classification data were exported from QuPath in Excel format and imported into Python for downstream analysis. Cell coordinates and classification labels were extracted and translated into defined cellular phenotypes via a pre-specified rules table (curated CSV). Here, we defined a proliferative tumor subphenotype as cells with Ki-67 positive, negative for all other markers in the panel. Spatial analysis was performed using the SciMap framework. aceted boxplots illustrating the distribution of average shortest distances from each phenotype to target phenotype.

#### Comparison of SCLC_TR to external signatures and datasets and tumor-reactive T cell prediction (ROC Analysis)

To benchmark the performance of the internally derived SCLC_TR signature, two published tumor-reactive gene signatures, NeoTCR8 and TR_30, were selected for comparative analysis. Each gene signature (SCLC_TR, NeoTCR8, and TR_30) was scored across all CD8⁺ T cells using the UCell scoring approach, implemented via the AddModuleScore_UCell function in the Seurat R package (v4.2.0). For visualization and comparison of expression patterns, signature scores were overlaid on UMAP embeddings and summarized in violin plots. Expression levels were compared across defined CD8⁺ T cell subsets, including both tumor-reactive and non-reactive populations, based on prior clonotype annotations. The classification performance of each signature was assessed using ROC curves and Area Under the Curve (AUC) metrics. ROC analysis was performed using the pROC R package (v1.18.0), with binary classification labels defined based on tumor-reactivity status (reactive vs. non-reactive) as determined through functional screening. Signature scores served as the predictor variable. AUC values were calculated using the roc() function, and comparisons across signatures were conducted within the same dataset to evaluate discriminatory power. To assess cross-context applicability, the SCLC_TR signature was applied to an external single-cell dataset derived from pancreatic ductal adenocarcinoma patients, as published by Meng et al.^29^ Pre-processed expression data and T cell annotations for the PDAC dataset were obtained from publicly available sources (NCBI BioProject PRJNA914983). Signature scores were computed using the same scoring and classification framework as described above to evaluate generalizability across distinct TMEs. For the prediction of tumor-reactive CD8+ T cells, the score for SCLC_TR signature served as the predictor variable. AUC values were calculated using the roc() function. The Youden index was used to calculate the optimal cutoff values for TR_30 (cutoff = 0.22).

#### Survival analysis using bulk RNA-seq

To test clinical relevance, the SCLC_TR signature was also evaluated in a bulk RNA-seq dataset from a previously published SCLC patient cohort (George et al., Nature 2015). Gene expression matrices and clinical survival data were obtained from the corresponding repository. 72 patients were stratified into high and low SCLC_TR signature score groups using median-based thresholding. Survival curves were generated using the survival (v3.3-1) and survminer (v0.4.9) R packages. Log-rank tests were used to assess statistical differences between groups, and hazard ratios were calculated using Cox proportional hazards models.

#### Signature gene overlap and expression analysis

Gene overlap among the SCLC_TR, NeoTCR8, and TR_30 signatures was visualized using Venn diagrams, generated via the VennDiagram R package (v1.7.3). Shared and unique gene features were annotated, and selected genes of interest were further analyzed for expression across all CD8⁺ T cell subsets using violin plots and heatmaps. Genes consistently found in all 3 signatures were selected for deeper exploration, with expression levels assessed across both reactive and non-reactive T cell populations in the SCLC dataset.

#### Antigen specificity prediction and TCR ranking based on SCLC_TR signature

TCR clonotypes with predicted viral, human and plant-derived antigen specificities were identified by VDJmatch and filtered for db confidence >= 1. TCR clonotypes were ranked by UCell scoring based on our newly identified SCLC_TR signature. Top TCR candidates with the highest maximum score for each clonotype besides the already validated ones were selected for further validation via ELISpot as described above.

