## Supplementary Figures and Table for "Mapping the immune landscape in small cell lung cancer unveils a distinct tumor-reactive CD8+ T cell molecular signature"

Overview

Supplementary Figure 1. Immune cell subset identification with myeloid cells at higher resolution

Supplementary Figure 2. Inter-tumor and intra-patient insights via scRNA seq analysis

Supplementary Figure 3. SCLC cell lines harbor a high mutation burden, closely resembling the parent tumors

Supplementary Figure 4: Expansion of SCLC TILs generates a tumor-specific immune response, enriching CD8+ T cells and tumor-reactive TCRs

Supplementary Figure 5: Characterisation of viral TCRs, tumor spheroid disintegration by tumor-reactive T cells and tumor-reactive TCRs distribution

Supplementary Figure 6: Enrichment and prediction of tumor-reactive and non-tumor reactive CD8+ T cells and functional implications in SCLC TME

Supplementary Figure 7: Overlap of SCLC_TR signature genes with two external signatures

Supplementary Table 1: Patient cohort, sample information, and post-filtering scRNA-seq/scTCR-seq sample quality-control metrics

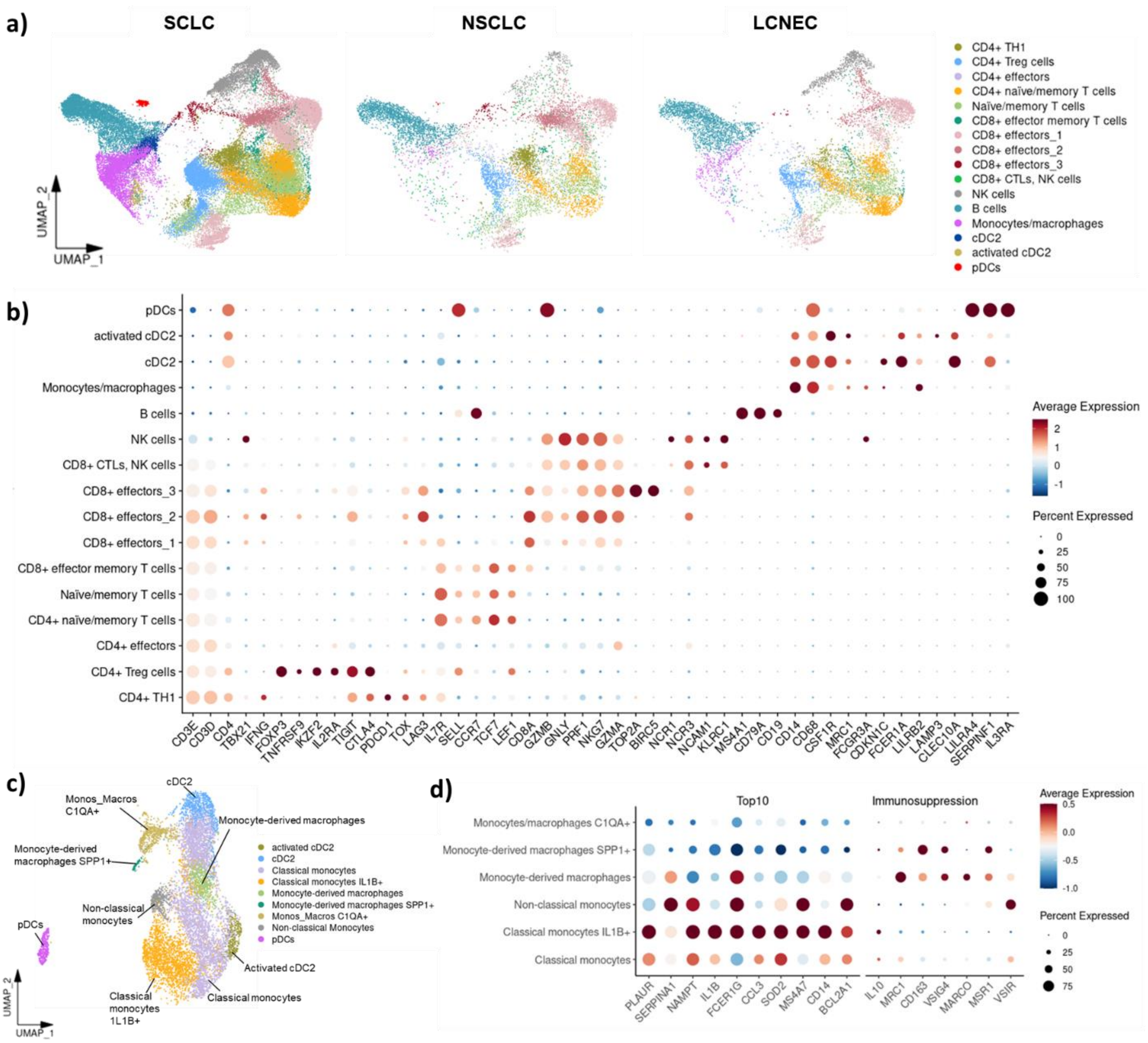


**Supplementary Figure 1. Immune cell subset identification with myeloid cells at higher resolution**

(a) UMAP (from main Fig. 1a) split according to tumor biopsies of SCLC, NSCLC and LCNEC, respectively, is depicted. Each dot on the UMAP represents a single cell and 16 broad immune cell subsets are annotated and marked by colour code. (b) Dot plot demonstrates expression of canonical cell markers in respective immune cell subsets shown in Fig 1b. Legend on the right signifies expression score and the size of the dot shows the expression by percentage of the population. (c) UMAP of subclustered myeloid cells derived from the UMAP in Fig. 1b shows identified additional clusters constituting the monocytes/macrophages cluster. (d) Dot plots show the top 10 differentially expressed genes between Monocytes/macrophages derived from SCLC compared with NSCLC and LCNEC plotted for all subclustered myeloid cells (adjusted $p < 0.05$, $\log_2$ fold change > 0.5, left side) and selected immunosuppressive markers for the same groups (right side). Color intensity represents scaled average gene expression, and dot size indicates the percentage of cells expressing each gene.

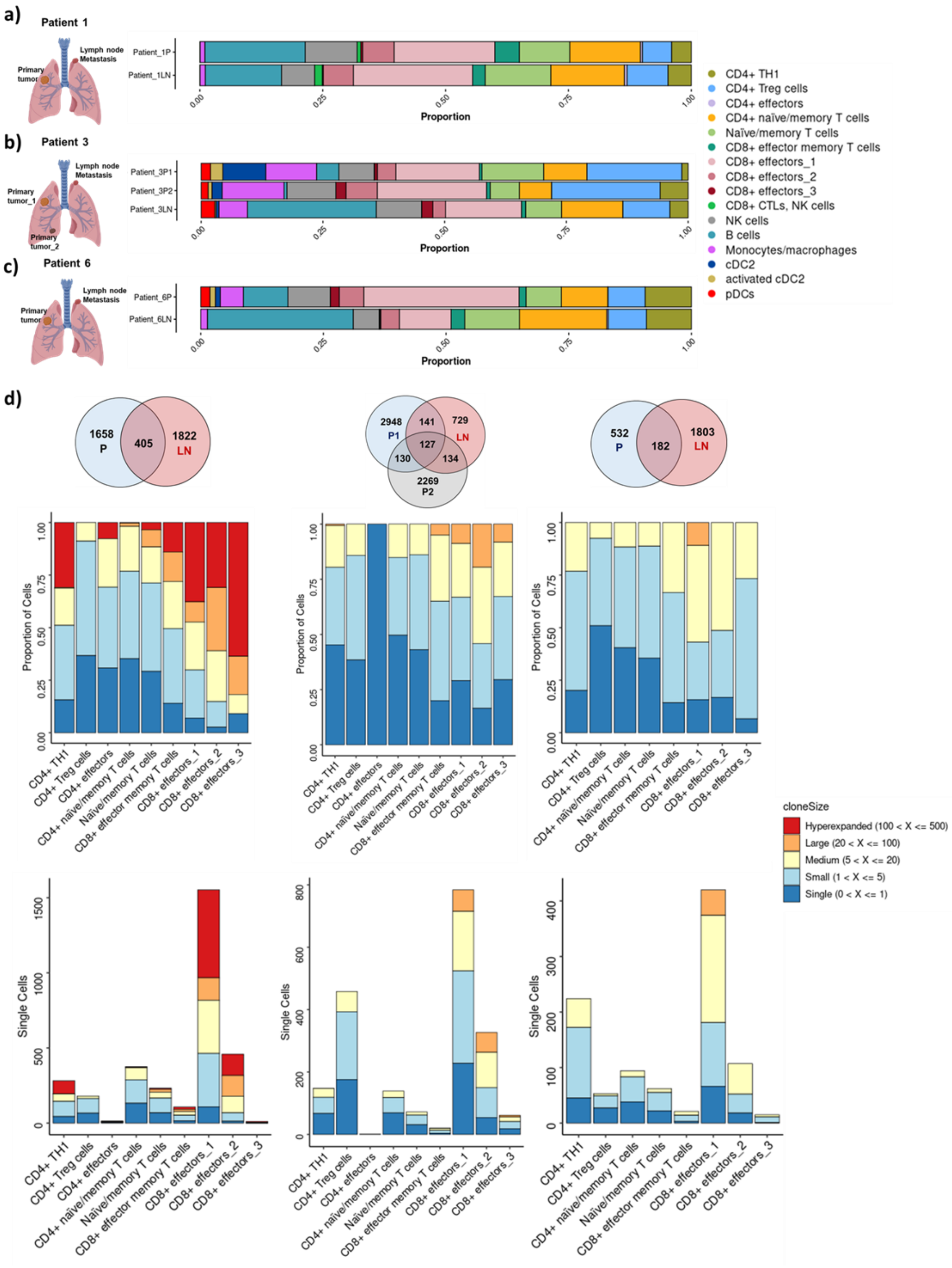


**Supplementary Figure 2. Inter-tumor and intra-patient insights via scRNA seq analysis**

(a-c) Schematic presentation of samples collected from patient 1 (a), patient 3 (b) and patient 6 (c). The bar plots illustrate the distribution of immune cell proportions in the primary tumor (P) and lymph node metastasis (L) of the patients 1 and 6 (a,c) and the distribution of immune cell proportions in the primary tumor (P1), primary tumor_2 (P2) and lymph node metastasis (L) of patient 3 (c), respectively. Image created with BioRender.com. (d) The Venn diagrams show overlapping and unique clones between primary region (P) and lymph node metastasis

(LN) in patient 1, 6 and 3, respectively. The bar plot below represents the phenotype of the cells presented by the shared clones across different sites in proportions (upper row) and absolute numbers (lower row), respectively. Each bar also contains the proportion of the clone size with the legend on the right side representing classification of the different cell numbers per clonal expansion compartment. The legend on the right side is consistent for all the bar plots.

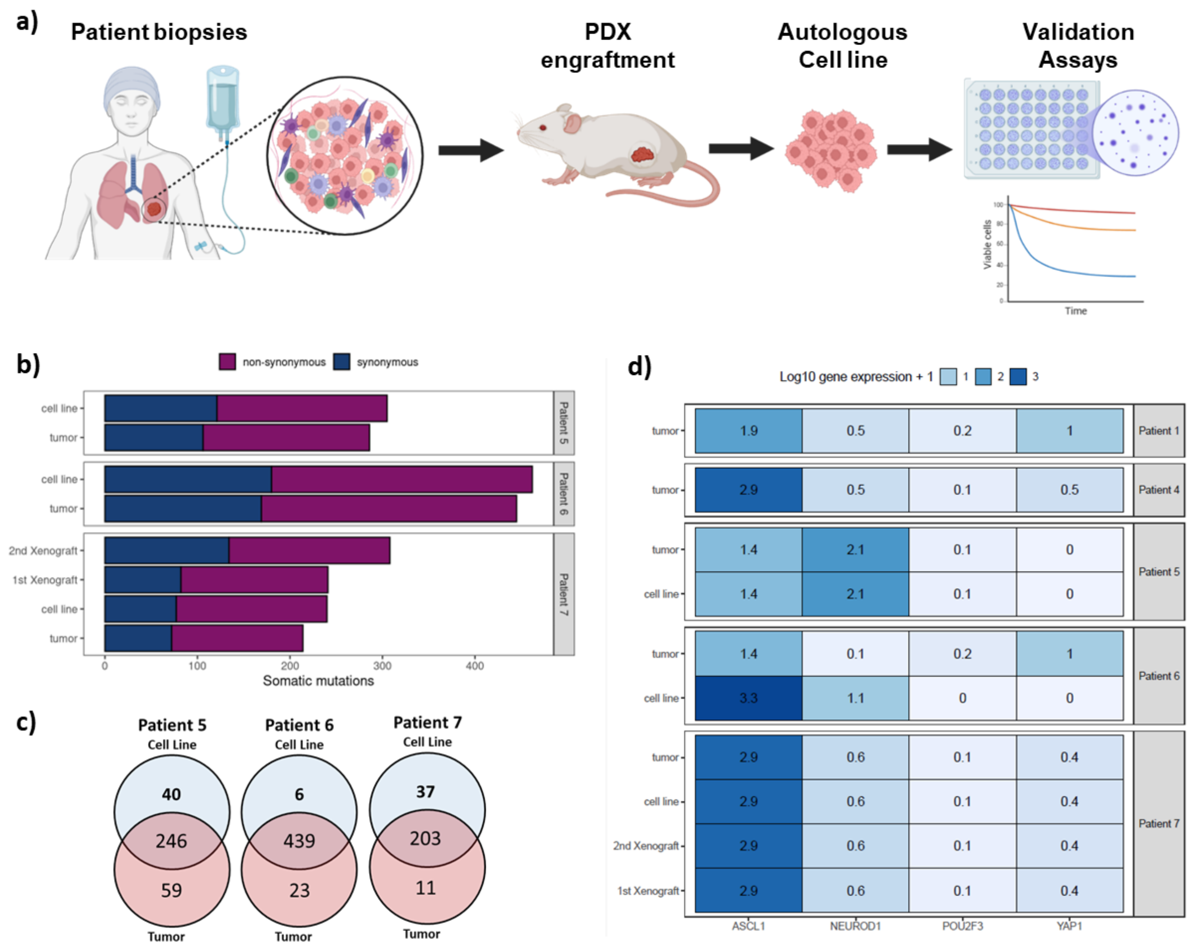


**Supplementary Figure 3. SCLC cell lines harbor a high mutation burden, closely resembling the parent tumors**

(a) Graphical summary of the experimental design for PDX engraftment and autologous cell line generation from biopsies from SCLC patient 5, 6 and 7 for subsequent use in downstream validation assays. Image created with BioRender.com. (b) Number of somatic mutations detected in the tumor and patient derived xenografts or cell lines from SCLC patient 5, 6 and 7. (c) The Venn diagrams show mutation overlap between cell lines and tumor biopsies of patient 5, 6 and 7. (d) The heatmap demonstrates expression of respective subtype markers ASCL1, NEUROD1, POU2F3 and YAP1 ASCL1, NEUROD1, POU2F3 and YAP1, shown in a heatmap for tumor biopsies and where applicable patient cell lines from patient 1, 4, 5, 6, and 7.

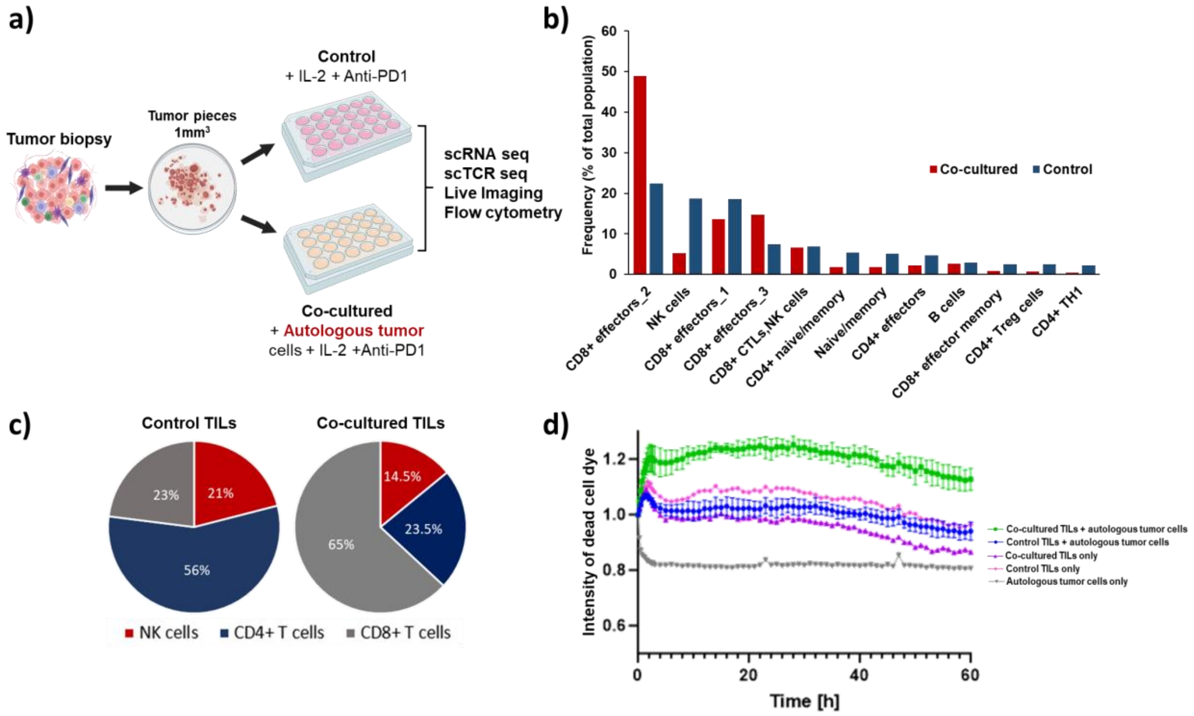


**Supplementary Figure 4: Expansion of SCLC TILs generates a tumor-specific immune response, enriching CD8+ T cells and tumor-reactive TCRs**

(a) Experimental setup for TIL expansion from tumor biopsy of Patient 6 is shown. For the ‘Control’ group tumor cells were expanded in the presence of IL-2 (100U/mL, for T cell proliferation) and anti-PD1, to counteract the induction of PD-L1, whereas for ‘Co-cultured’ group autologous tumor cells were additionally added to the culture. Image created with BioRender.com. (b) Results from scRNA-seq data illustrating distribution of cell subset proportions between the two groups ‘Control’ and ‘Co-cultured’. (c) Flow cytometric indicating a 6% reduction in NK cells and a 42% increase in CD8+ T cells in the "Co-cultured" TILs compared to the "Control" group. (d) Results from CQ1 tumor killing assay post-expansion of the conditions ‘Control’ vs ‘Co-cultured’ are depicted. TILs co-cultured with tumor cells (green) demonstrated significantly enhanced killing potential compared to those that were not exposed to tumor cells during co-culture (blue).

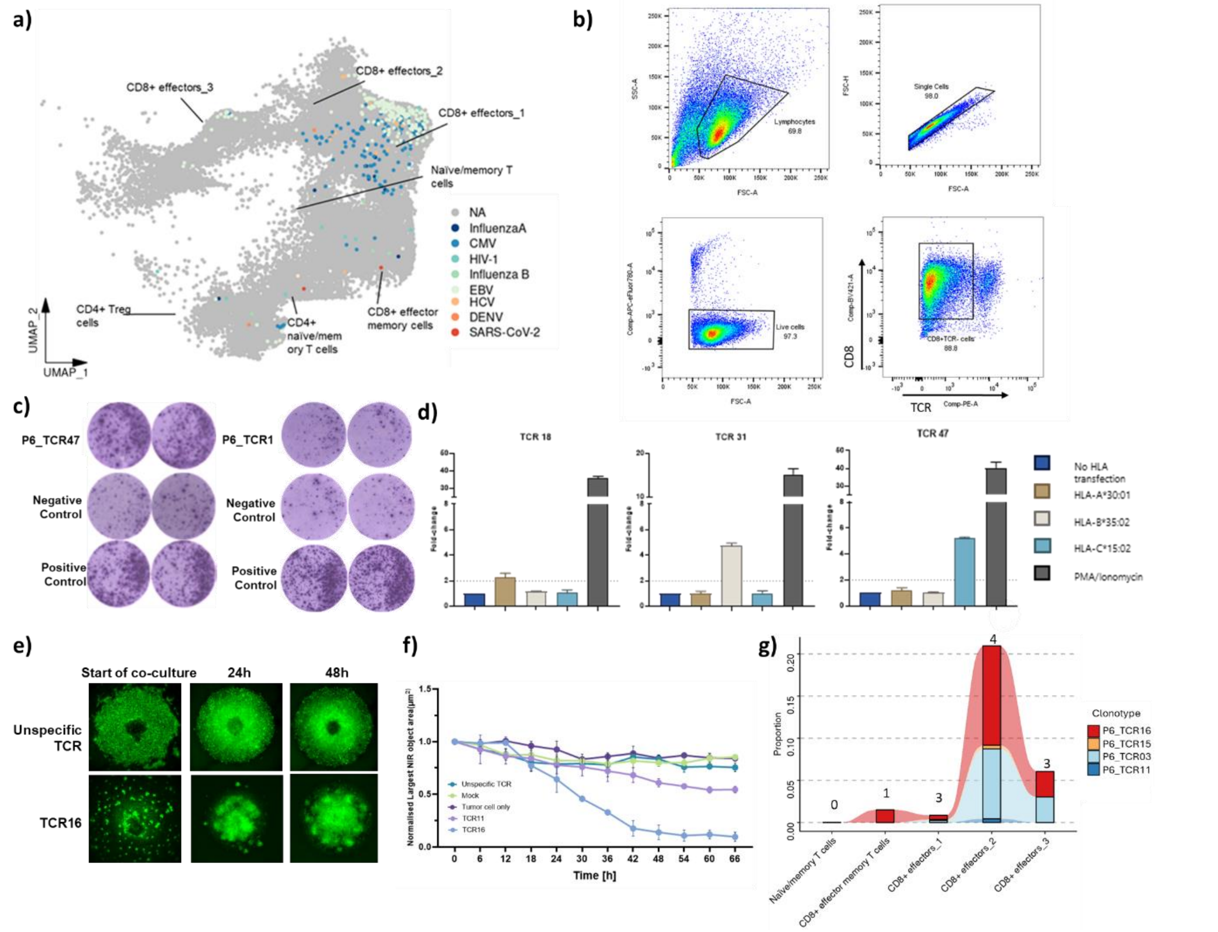


**Supplementary Figure 5: Characterisation of viral TCRs, tumor spheroid disintegration by tumor-reactive T cells and tumor-reactive TCRs distribution**

(a) The UMAP of subsetted T cells represents the viral TCRs identified and excluded from the validation pipeline to find tumor-specific TCRs. (b) The flow cytometry results show the gating strategy and final dot plots for TCR and CD8 expression. (c) CTL ImmunoSpot images compare a reactive TCR (TCR47) and a non-reactive TCR (TCR1). (d) Reactivity screen using transgenic Jurkat-NFAT reporter cells shows the reactivity of individual TCRs towards K562 cells as orthogonal APC expressing the respective HLA molecule as indicated. (e) Images from the 3D tumor spheroid killing assay for TCR16 from patient 6, where T cells are stained in green, demonstrate clustering of T cells upon tumor spheroid attack as shown in Fig. 5f, when tumor-reactive TCR-expressing T cells are introduced (lower panel), compared to control T cells with non-specific TCRs (upper panel). (f) The 3D tumor-killing assay for two TCRs from patient 5 shows normalized spheroid intensity (y-axis) over time (x-axis), with each TCR represented. (g) The changing frequencies of tumor-reactive TCRs from the primary tumor of patient 6 across different T cell subsets from the SCLC microenvironment are depicted. The y-axis indicates the proportion of respective tumor-reactive TCR. The number on top of each bar represents the number of TCRs traced in each condition.

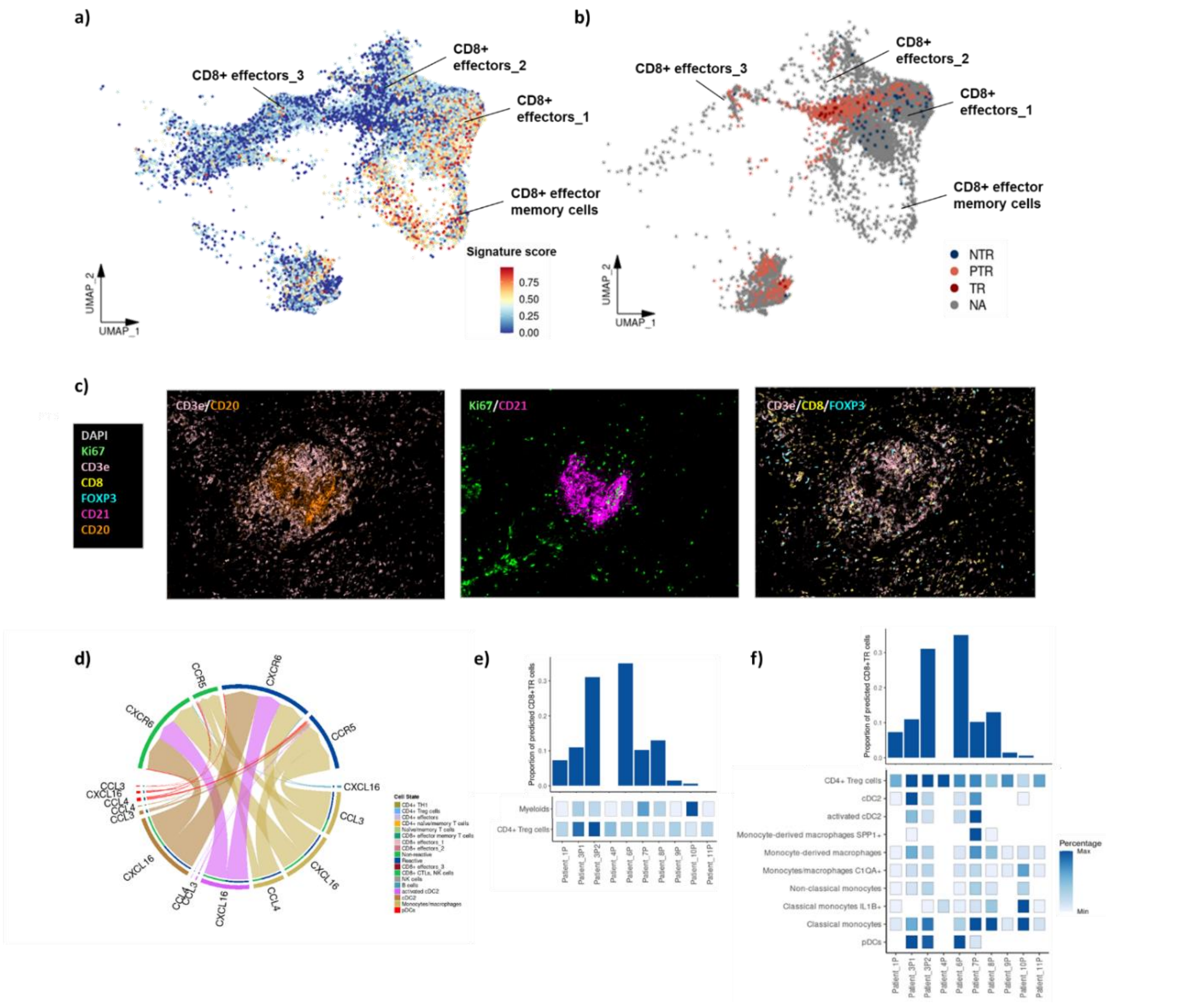


**Supplementary Figure 6: Enrichment and prediction of tumor-reactive and non-tumor reactive CD8+ T cells and functional implications in SCLC TME**

(a) The feature plot shows the spatial distribution of CD8+ T cells from SCLC patient biopsies enriched in the molecular signature of non-reactive T cells (SCLC_NTR). (b) UMAP visualization shows confirmed tumor-reactive (TR), confirmed non-tumor-reactive (NTR), and predicted tumor-reactive (PTR) clonotypes based on ROC analysis. (c) Representative high-plex immunohistochemistry (IHC) staining, showing 2-3 markers at a time, illustrates the spatial distribution of mature TLS structures within a SCLC tumor sample with a magnified view of a representative TLS characterized by a core of B cells (CD20+) and follicular dendritic cells (CD21+) surrounded by a ring of CD4+ $T_{reg}$ cells (CD3+ FOXP3+ CD8-), T helper cells (CD3+ FOXP3- CD8-) and cytotoxic CD8+ T cells (CD3+ CD8+) in a peritumoral region (Ki67+) (scale bar: 1.5 mm; inset scalebar: 50 µm). (d) Cell-cell communication analysis showcases elevated predicted chemokine and cytokine signaling between reactive and non-reactive T cells on the top as target cells and pDCs, monocytes/macrophages, activated cDC2, cDC2, and B cells on the bottom as source cells in all SCLC patients. (e-f) The bar plots illustrate the proportion of predicted tumor-reactive CD8+ T cells within the primary tumor region for each SCLC patient. The heat maps below represent the proportion of corresponding immunosuppressive cell subsets in the total tumor microenvironment for each primary tumor region. The gradient scale on the right transitions from white to dark blue, indicating increasing proportions.

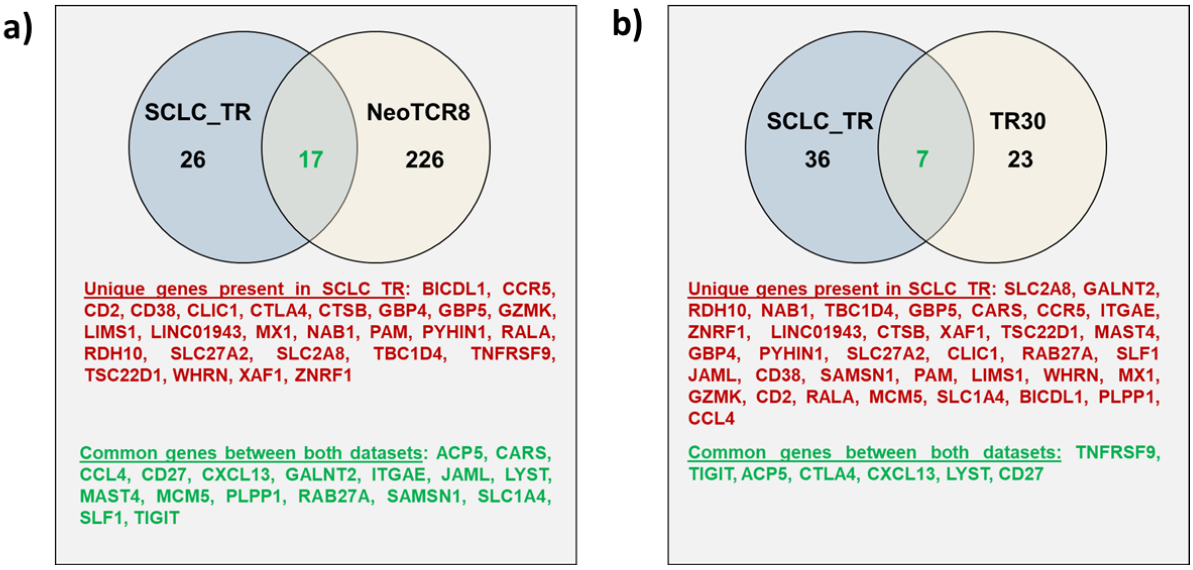


**Supplementary Figure 7: Overlap of SCLC_TR signature genes with two external signatures**

(a-b) Overlapping markers between SCLC TR signature comprising of 43 genes and the NeoTCR8 signature comprising of 243 genes (a) and the TR_30 signature comprising of 30 genes (b)

**Supplementary Table 1: Patient cohort, sample information, and post-filtering scRNA-seq/scTCR-seq sample quality-control metrics**

| Patient_ID | Sample_backgrou | Group | cell_number | TCR_clones | feat_length | feat_max | feat_median | feat_mad | feat_upper_cutoff | mt_length | mt_max | mt_median | mt_mad | mt_upper_cutoff |
|---|---|---|---|---|---|---|---|---|---|---|---|---|---|---|
| Patient_1P | Primary | Day0_SCLC | 5720 | 1964 | 6795 | 11200 | 2021 | 421,06 | 3284,18 | 6795 | 70,01 | 3,19 | 1,15 | 6,64 |
| Patient_1LN | Lymph node meta | Day0_SCLC | 4559 | 1792 | 5607 | 9085 | 2585 | 532,25 | 4181,76 | 5607 | 71,37 | 4,16 | 1,6 | 8,96 |
| Patient_2LN | Lymph node meta | Day0_SCLC | 1290 | 510 | 3024 | 6732 | 1391 | 871,77 | 4006,31 | 3024 | 75,67 | 6,11 | 5,81 | 23,55 |
| Patient_3LN | Lymph node meta | Day0_SCLC | 4957 | 1768 | 8677 | 8109 | 1554 | 885,11 | 4209,34 | 8677 | 78,59 | 5,3 | 3,64 | 16,21 |
| Patient_3P1 | Primary | Day0_SCLC | 6283 | 2246 | 11983 | 7418 | 1208 | 928,11 | 3992,32 | 11983 | 83,14 | 6,65 | 5,32 | 22,62 |
| Patient_3P2 | Secondary metast | Day0_SCLC | 2398 | 922 | 4220 | 7338 | 1506 | 1136,41 | 4915,24 | 4220 | 75,43 | 5,84 | 5,19 | 21,42 |
| Patient_4P | Primary | Day0_SCLC | 104 | 74 | 205 | 1871 | 588 | 200,15 | 1188,45 | 205 | 32,11 | 4,21 | 1,74 | 9,42 |
| Patient_5Be | PBMCs cocultured | PBMC | 425 | 354 | 639 | 6971 | 1978 | 665,69 | 3975,06 | 639 | 27,11 | 4,15 | 2,18 | 10,68 |
| Patient_5Te | Expanded TILs | Expanded_SCLC | 2400 | 308 | 3285 | 4873 | 2223 | 560,42 | 3904,27 | 3285 | 57,97 | 4,03 | 1,3 | 7,91 |
| Patient_6P | Primary | Day0_SCLC | 1981 | 764 | 5087 | 7516 | 2511 | 597,49 | 4303,46 | 5087 | 75,39 | 3,85 | 1,55 | 8,51 |
| Patient_6LN | Primary | Day0_SCLC | 4149 | 1933 | 3640 | 9300 | 1909 | 899,94 | 4608,81 | 3640 | 82,44 | 3,25 | 2,09 | 9,53 |
| Patient_6LNe | Expanded TILs | Expanded_SCLC | 5597 | 1248 | 1766 | 8174 | 1926 | 942,19 | 4752,58 | 1766 | 73,28 | 3,97 | 2,67 | 11,97 |
| Patient_6Pe_Contr | Expanded TILs | Expanded_SCLC | 5445 | 946 | 7020 | 9430 | 2732,5 | 1065,25 | 5928,24 | 7020 | 61,05 | 7,87 | 1,84 | 13,4 |
| Patient_6Pe_Cocu | Expanded TILs | Expanded_SCLC | 7067 | 332 | 1451 | 6078 | 1528 | 361,75 | 2613,26 | 1451 | 47,06 | 2,64 | 1,01 | 5,68 |
| Patient_7P | Primary | Day0_SCLC | 1218 | 375 | 3403 | 8905 | 2232 | 683,48 | 4282,44 | 3403 | 55,35 | 2,36 | 0,91 | 5,08 |
| Patient_7Pe | Expanded TILs | Expanded_SCLC | 1156 | 673 | 1406 | 9383 | 1984,5 | 420,32 | 3245,45 | 1406 | 78,34 | 2,71 | 1,05 | 5,85 |
| Patient_8P | Primary | Day0_SCLC | 2634 | 1217 | 5192 | 7398 | 2084 | 793,19 | 4463,57 | 5192 | 71,42 | 1,86 | 1,02 | 4,92 |
| Patient_9P | Primary | Day0_SCLC | 1050 | 752 | 3054 | 6632 | 1851 | 399,56 | 3049,68 | 3054 | 51,57 | 1,76 | 0,72 | 3,94 |
| Patient_10P | Primary | Day0_SCLC | 3388 | 546 | 9054 | 9007 | 2495 | 1153,46 | 5955,39 | 9054 | 88,41 | 7,62 | 4,45 | 20,97 |
| Patient_11P | Primary | Day0_SCLC | 2484 | 1885 | 9881 | 7482 | 2727 | 855,46 | 5293,38 | 9881 | 79,41 | 5,31 | 1,59 | 10,09 |
| Patient_12P | Primary | Day0_NSCLC | 6063 | 2148 | 13200 | 7505 | 1905,5 | 1690,91 | 6978,22 | 13200 | 81,24 | 8,74 | 8,36 | 33,82 |
| Patient_13P | Primary | Day0_NSCLC | 1629 | 54 | 3370 | 2809 | 465 | 232,77 | 1163,3 | 3370 | 53,73 | 8,11 | 3,94 | 19,93 |
| Patient_14P | Primary | Day0_NSCLC | 8210 | 2581 | 11812 | 7481 | 1886 | 610,09 | 3716,27 | 11812 | 63,31 | 4,62 | 2,07 | 10,82 |
| Patient_15P | Primary | Day0_LargeCellNE | 2878 | 1475 | 3960 | 7122 | 2045,5 | 543,37 | 3675,62 | 3960 | 54,59 | 2,31 | 1,05 | 5,46 |
| Patient_16P | Primary | Day0_LargeCellNE | 2934 | 1254 | 3819 | 7286 | 1902 | 535,22 | 3507,66 | 3819 | 69,46 | 3,21 | 1,2 | 6,83 |
| Patient_17P | Primary | Day0_LargeCellNE | 3335 | 2075 | 4535 | 7039 | 1643 | 376,58 | 2772,74 | 4535 | 31,05 | 2,78 | 1,06 | 5,97 |
